# Retrieving high-resolution chromatin interactions and decoding enhancer regulatory potential *in silico*

**DOI:** 10.1101/2020.11.10.376533

**Authors:** Eduardo Gade Gusmao, Athanasia Mizi, Lilija Brant, Argyris Papantonis

**Affiliations:** Institute of Pathology, University Medical Center Göttingen, 37075 Göttingen, Germany; Center for Molecular Medicine Cologne, University of Cologne, 50931 Cologne, Germany

**Author notes:** Correspondence to: A.P.

**Keywords:** chromatin loops, Hi-C, Dirichlet processes, mixture models, *cis*-element, topological domain

## Abstract

The advent of the chromosome conformation capture (3C) and related technologies has profoundly renewed our understaning of three-dimensional chromatin organization in mammalian nuclei. Alongside these experimental approaches, numerous computational tools for handling, normalizing, visualizing, and ultimately detecting interactions in 3C-type datasets are being developed. Here, we present *Bloom*, a comprehensive method for the analysis of 3C-type data matrices on the basis of Dirichlet process mixture models that addresses two important open issues. First, it retrieves occult interaction patterns from sparse data, like those derived from single-cell Hi-C experiments; thus, *bloomed* sparse data can now be used to study interaction landscapes at sub-kbp resolution. Second, it detects enhancer-promoter interactions with high sensitivity and inherently assigns an interaction frequency score (IFS) to each contact. Using enhancer perturbation data of different throughput, we show that IFS accurately quantifies the regulatory influence of each enhancer on its target promoter. As a result, *Bloom* allows decoding of complex regulatory landscapes by generating functionally-relevant enhancer atlases solely on the basis of 3C-type of data.

## Introduction

Mammalian chromatin is a long biopolymer that must fit the confines of μm-wide cell nuclei, hence the multiple layers of folding applied across its length by converging and conflicting forces.^1^ Despite its packing, chromatin needs to fulfill its regulatory roles — i.e., ensure cell type-specific gene expression programs and faithful cell cycle progression.^2^ The advent of chromosome conformation capture (3C) techniques over the last decade has allowed us to map chromatin folding at increasing resolution and throughput,^3^ and thus profoundly renewed our view of how dynamic and static chromatin configurations inform such cell type-specific regulation.

Mammalian chromosomes occupy individual territories, and each chromosome can be divided into Mbp-long “A” (mostly transcriptionally-active) and “B” (mostly repressed) compartments that tend to homotypically interact.^4^ A-A/B-B interactions are counteracted by the formation of sub-Mbp topologically-associating domains^5^ that mix and insulate chromatin such that intraTAD interactions are favoured over interTAD ones. At a finer scale, multi-kbp-long loops of different types have been discovered. Strong “structural” loops anchored at convergent CTCF-bound sites give rise to insulated “contact domains” that rely on cohesin complexes for their stabilization.^4,6^ Within these domains, the directed interplay between enhancers and gene promoters fine-tunes gene expression, although such regulatory interactions have now been recorded over larger scales too.^7–9^

These spatial configurations reflect a structure-to-function relationship that spans orders of magnitude in scale and that still requires elucidation. For instance, it is pertinent to gain a high-resolution understanding of single-cell chromatin folding and to obtain quantitative information on gene-regulatory looping. The former would allow us to address cell-to-cell heterogeneity, but current approaches are hampered by contact sparsity and low resolution.^10^ The latter would help us stratify *cis*-element interactions based on their actual regulatory impact, but it remains challenging at the whole-genome level as it requires sub-kbp-resolution Hi-C data, different epigenomic information for mapping enhancers (e.g., TF positioning, histone and DNA modifications, chromatin accessibility), as well as laborious assays to functionally test individual enhancers (e.g., CRISPR-based assays).^11,12^ Resolving these issues can profoundly change our appreciation of 3D regulatory wiring along mammalian chromosomes and provide a gateway into dissecting diseases stemming from alterations in chromatin folding or enhancer variation.^13^

Motivated by these unresolved issues, we developed a novel *in silico* approach called *Bloom* to (i) assuage the sparsity of contact maps derived from 3C-type data; (ii) reduce saturation and contact overrepresentation in dense maps; (iii) score interactions in a quantitative and interpretable manner. *Bloom* inherently derives a non-distribution-bound metric that reflects the actual regulatory strength of enhancer-promoter contacts. Thus, end-users can readily deduce genome-wide enhancer atlases that are simultaneously informative of their regulatory potential and virtually independent of initial data sparsity or structure from Hi-C maps alone.

## Results

### The workflow behind Bloom

Alongside 3C-based technologies, computational tools also represent a particularly active research area. Thus, for detecting, assessing, and processing loop-level/subTAD interactions, multiple tools have been developed to date (for a comparison see ref. 14). Of course, these are not without caveats and are generally incapable of dealing with sparsely-populated contact matrices. This was our initial motivation for developing *Bloom*. *Bloom* uses a raw (unprocessed) chromatin contact matrix as input. Such a matrix can be derived from Hi-C (“all-to-all” contact mapping), capture-based approaches like CaptureC, T2C or Promoter Capture Hi-C (“many-to-many” with probe subselection) or immunoselection-based assays like HiChIP or PLAC-seq (“many-to-many” based on a specific chromatin-associated factor).^3^ The only constraint is input of a matrix with an equal number of columns and rows in any standard 3C-data format^15,16^ as *Bloom* handles them as a simple indexed list and, thus, turns the information retrieval computational complexity to zero.

The core computational workflow of *Bloom* (see **Fig. 1a**) is comprised of three main steps. First, an initial check is used to verify whether the matrix is sparse or dense and informative data components are next extracted — i.e., the most independent rows/columns from one another. Dense matrices are subjected to the established “independent component analysis” (ICA),^17^ while sparse ones undergo a novel type of processing called “sparse independent component analysis” (SICA) to recover independent components (see **Methods** for details). In a second step, experimentally-derived biases (e.g., self-ligations) are removed with the intent to ease computation by reducing the norm of the contact matrix. This takes place via a new and fast matrix-balancing algorithm based on Osborne matrix balancing^18^ that we call “generalized osborne-balancing algorithm” (GOBA). It is important to point out that our matrix factorizations (ICA/SICA) are not used for dimensionality reduction; instead, they indicate important rows/columns for “balancing” by the ensuing GOBA approach. In a third and last step, an “iterative hierarchical Dirichlet process mixture model” (iDPMM) is applied.^19^ This aims at unveiling occult contact patterns in the matrix. Briefly, the “balanced” contact matrix is fit to multiple Dirichlet distribution instances. Importantly, the statistical distributions that the Dirichlet process consists of closely follow inherent data distribution — i.e., our approach does require fitting data to known distributions. Next, an iterative phase takes place where, at each iteration, novel data is (i) imputed based on information from the previous iterations and (ii) corrected by re-evaluating parameters for each individual distribution. iDPMM is a non-binning-specific method — i.e., contact map resolution can be set to any user-defined target (even down to “nucleosome-resolution” like that afforded by Micro-C^20,21^). To avoid the computational burden of adding a “convergence” check at every iteration — i.e., a stop point to check if no significant information gain is to occur — we perform “convergence verification” after a defined number of iterations (empirically defined at 50 iterations for all data and cell types tested in this study). The key aspect of iDPMM is that the strength of each chromosomal contact is automatically reported through convolution of Dirichlet instances as a unified “interaction frequency score” (IFS). This renders the accuracy of subsequent analyses, such as compartment and TAD detection more robust.

**Figure 1.**
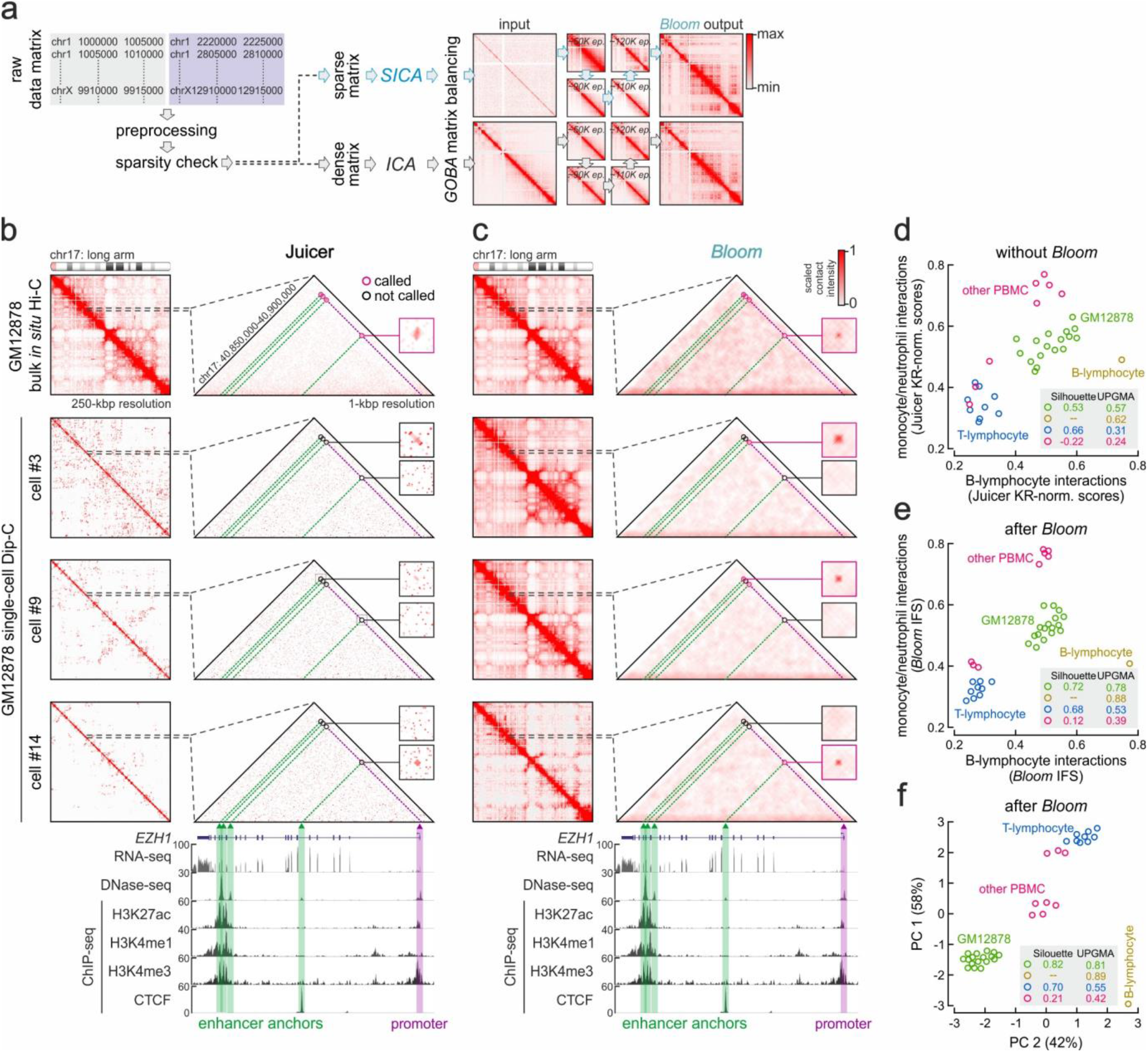
*Bloom* retrieves rich interaction profiles from single-cell Hi-C data. **(a)** Overview of the *Bloom* workflow. **(b)** Bulk *in situ* Hi-C (*top*) and single-cell Dip-C data from three randomly-selected GM12878 cells (#3, #9 and #14) analyzed using Juicer. *Zoom in*: interactions in the *EZH1* locus (*triangles*) aligned to ENCODE RNA-/ChIP-seq data and enhancer-promoter contacts identified (*circles*). **(c)** As in panel b, but analyzed using *Bloom*. **(d)** Scatter plot stratifying PBMC and GM12878 populations based on Dip-C data analyzed using Juicer. Silhouette and UPGMA scores for each cell population are shown (*inset*). **(e)** As in panel d, but using data analyzed using *Bloom*. (f) As in panel e, but using PCA.

### Bloom retrieves high-resolution contact matrices from single-cell Hi-C data

To test the capacity of *Bloom* for retrieving meaningful high-resolution contact matrices from sparse Hi-C data, we turned to single-cell Hi-C experiments that are in inherently sparse and heretogeneous. We used Dip-C data generated on single lymphoblast cells (GM12878),^22^ as well as bulk kbp-resolution *in situ* Hi-C data from the same cell type as reference.^4^ All data were processed and visualized in parallel via the standard Juicer/HiCCUPS combo^15^ or via *Bloom*. Using the highly expressed *EZH1* locus on chr17 as a representative example, we compared contact maps at low and high resolution. At 250 kbp-resolution, *Bloom* transformed the sparse Dip-C contact matrices of the whole chr17 long arm from three randomly-selected cells (#3, #9, and #14 in ref. 22) into detailed maps closely resembling the population reference (**Fig. 1b,c**). Despite subtle differences in compartmentalization strength, *Bloom* retrieved very similar contact architectures for the three individual cells.

Zooming into the *EZH1* domain at 1-kbp resolution, *in situ* Hi-C data contain interactions connecting the *EZH1* promoter with three clustered intragenic enhancers and a mid-gene positioned CTCF site. These contacts were still detected after applying *Bloom* to the bulk data (suggesting that *Bloom* does not convolute high-resolution data such that disciriminatory power is lost; **Fig. 1b,c**). However, none of these interactions could be detected after standard processing of Dip-C data (**Fig. 1b**). In contrast, *Bloom*, unveiled different contacts in each cell (**Fig. 1c**). This is in line with the heterogeneity expected of single-cell chromatin folding, but may also explain the variability in EZH1-target repression that was reported for lymphoblasts.^23^

Nonetheless, to ensure that such differentially-observed contacts are not random effects, and to exclude early or late convergence to the same output despite different input data (a common issue with approaches using unsupervised stochastic learning, like *Bloom*), we present an additional example in the immunoglobulin light chain *λ* locus. Here, the 250 kbp-resolution maps of the long arm of chr22 again display contact patterns that closely match those of *in situ* Hi-C (**Fig. S1a,b**), but differences emerge in the 0.5 kbp-resolution matrices around the *IGLJ2*, *IGLJ3*, *IGLC3* and *IGLC6* gene cluster that is active in GM12878 cells. Several putative enhancers, all marked by H3K4me1, H3K27ac and DNase I hypersensitivity regions, are found downstream of these genes. The high activity of this locus yields particularly strong Dip-C signal and, thus, even standard analysis picks up enhancer-promoter contacts (**Fig. S1a**). These are recapitulated by *Bloom*, which however ignores some (in cells #3 and #9) or reveals others *de novo* (in cells #9 and #14; **Fig. S1b**). Although none of these contacts have been orthogonally validated, they agree with the differential expression of light chain genes in lymphoblasts. Together, these data confirm that *Bloom* does not obscure interactions in raw matrices, while also unveiling contact heterogeneity and not converging to a single folding state.

In order to validate that the contact profiles unveiled via *Bloom* are indeed occult in Dip-C data and not artefactual, we replicated the single-cell clustering analysis performed in the original publication.^22^ In brief, we calculated the average KR-normalized interaction scores in monocyte-/neutrophil-specific gene loci for each individual GM12878 or PBMC-derived cell (via Juicer) and plotted them against the same scores for lymphocyte-specific genes (Fig. 1d). In parallel, we repeated this analysis using the IFS values generated by *Bloom* (**Fig. 1e**). Comparison of the resulting plots shows better separation of cell types when using IFS, a fact corroborated by the calculation of “Silhouete” and UPGMA (unweighted pair group method with arithmetic mean) scores as metrics of intra-cluster conciseness and inter-cluster distances, respectively (see **Methods** for details). For example, the marked increase in GM12878 Silhouette score from 0.53 to 0.72 or T-lymphocyte UPGMA from 0.31 to 0.53 showcase *Bloom*‘s performance. Finally, principal component analysis of *Bloom*-derived values showed a homogeneous spread in variance, portraying higher dependence on this data type (PC1 and PC2 variabilities are quite similar at 58 and 42%; **Fig. 1f**). In summary, we show that *Bloom* reconstructs kbp-resolution occult cell type-specific information with high sensitivity and specificity without converging to a single result.

### Bloom enhances analysis of both sparse and dense contact maps

We recently introduced “intrinsic 3C” (i3C), a crosslinking-free method that allows interrogation of 3D chromatin interactions in nuclei under native conditions. i3C negates potential biases stemming from cell fixation and, thus, provided the first evidence for the topological restrictions imposed by TADs in a native context.^24^ i3C can map interactions with high signal-to-noise ratios that is advantageous when interrogating complex/dense landscapes.^25^ However, i3C contact matrices (from genome-wide iHi-C or capture-based iT2C experiments) suffer from sparsity, which renders their visual evaluation challenging.^24,26^ In addition, standard Hi-C analysis tools perform poorly on iHi-C data. The performance of *Bloom* on single-cell Dip-C data showed great promise for iHi-C data analysis too. To test this, we generated iHi-C using living mouse ES cell nuclei (mESCs; see **Methods**) for which high-quality matching *in situ* conventional Hi-C data is available.^27^ As before, raw iHi-C contact maps were essentially reduced to individual interaction points of high significance against very low background signal (**Fig. 2a**). This sparsity was alleviated by *Bloom* and the resulting contact maps now resembled those of *in situ* Hi-C, while also displaying fine interaction structure (**Fig. 2a**; despite differences in interaction depth, with >550 million in the conventional data versus ~78 million in iHi-C). We next used typical metrics, like compartment-level interactions and compartment insulation (at 100-kbp resolution) to show that *bloomed* iHi-C data show negligible (<4%) compartment differences compared to conventional mESC Hi-C (**Fig. 2b**), but stronger A/B compartment insulation (**Fig. 2c**). As regards TADs, rGMAP^28^ could not detect any in raw iHi-C, but in *bloomed* data >91% of TADs called were identical to those from conventional Hi-C (**Fig. 2d,e**). Again, insulation at *bloomed* iHi-C TAD boundaries was significantly stengthened (**Fig. 2f**) and the decay of contact frequencies with genomic distance was quite similar for the two methods (**Fig. 2g**).

**Figure 2.**
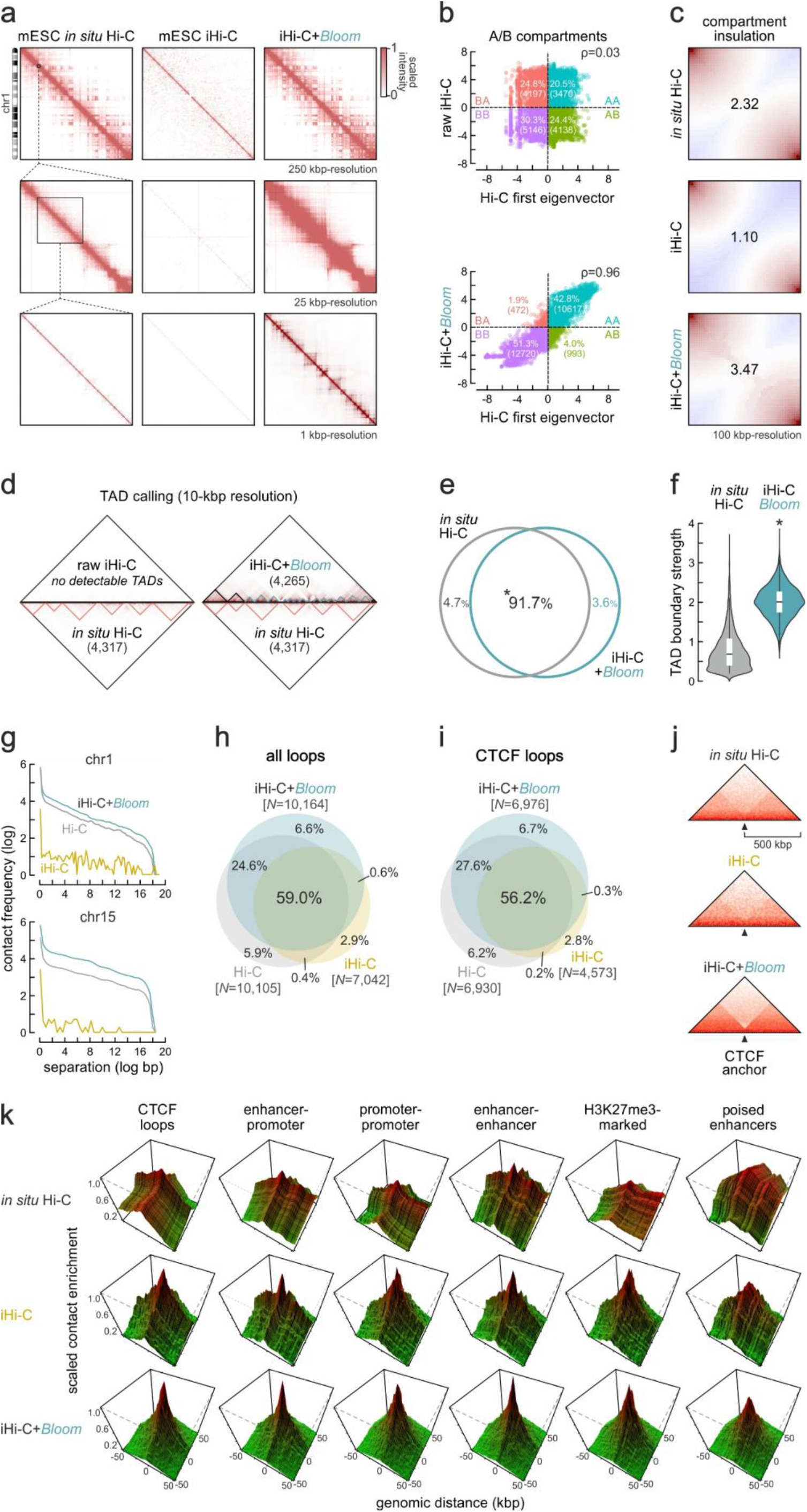
*Bloom* retrieves rich interaction profiles from sparse iHi-C data. (**a**) Exemplary contact maps from chr1 using *in situ* Hi-C (*left*), raw (*middle*) or *bloomed* iHi-C data from mESCs (*right*). (**b**) Scatter plots of A/B compartment value comparisons using data from panel a. (**c**) Saddle plots showing A-/B-compartment insulation in the data from panel a. (**d**) TADs in chr1 called using *in situ* Hi-C (*bottom triangles*) compared to those called using raw (*left*) or *bloomed* iHi-C data (*right*). TADs specific to *bloomed* iHi-C are denoted (*light blue triangles*). (**e**) Venn diagram showing the overlap between TADs called in Hi-C (*grey*) and *bloomed* iHi-C data (*light blue*). (**f**) Violin plots showing insulation strength at TAD boundaries using data from panel e. *: significantly different; *P*<10^−6^, Wilcoxon-Mann-Whitney test. (**g**) Line plots showing contact frequency decay with genomic distance for *in situ* Hi-C (*grey*), raw (*yellow*) or *bloomed* iHi-C (*light blue*). (**h**) Venn diagrams showing overlap of all loops called in data from panel a. *: more than expected by chance; *P*<10^−4^. (**i**) As in panel h, but for CTCF-anchored loops. (**j**) Heatmaps showing mean contact distribution in 1 Mbp around CTCF loop anchors from panel i. (**k**) 3D plots45 depicting Hi-C signal enrichment at CTCF-bound sites, active promoter-enhancer pairs, active promoters, active enhancers, H3K27me3-marked sites, and poised enhancers.

Next, we interrogated sub-Mbp loops and loop domains. Using HiCCUPS, we could call >10,000 loops in conventional Hi-C data, of which 6,930 were CTCF-anchored loops. In comparison, raw iHi-C yielded >7,000 loops with 4,573 being CTCF-anchored; in *bloomed* iHi-C these numbers increased to >10,150 and 6,976, respectively (**Fig. 2h**). 59 and 56% of all and of CTCF-anchored loops, respectively, were shared between the three datasets, but application of *Bloom* added a significant ~25% more loops to either overlap (**Fig. 2i**). Insulation calculated for CTCF loop anchors was also markedly enhanced following *blooming* of iHi-C (**Fig. 2j**). Subsequently, we stratified interactions based on their epigenetic demarcation. Looking into Hi-C versus iHi-C signal connecting active enhancers and promoters, we found that it was stronger for iHi-C and further enhanced following *Bloom* (**Fig. 2k**). Similar enhancement was obtained for all combinations tested, including contacts marked by H3K27me3 due to Polycomb-binding (**Fig. 2k**), shown to loop together poised promoters and/or enhancers in mESCs.^29^

To confirm *Bloom* performance on an independent dataset from a different cell type, we generated iHi-C data from iPSC-derived cardiomyocytes (CMs). Here, we used <1 million of these non-dividing cells via a modified iHi-C 2.0 protocol to minimize material losses.^26^ For comparison, we used *in situ* DNase Hi-C data generated on ~20 million CMs (with >5x more sequencing depth).^30^ Just like what we observed for mESC iHi-C, CM iHi-C 2.0 produced sparse matrices, but *Bloom* transformed these into detailed interaction maps closely resembling the conventional Hi-C profiles (**Fig. S2a**). This held true when interrogating compartments, TADs, and contact decay profiles (**Fig. S2b-g**). At the level of loops, raw iHi-C again performed well compared to conventional Hi-C, but *Bloom* markedly increased signal-to-noise ratios and contact discovery (**Fig. S2h-k**). Notably, applying *Bloom* to conventional mESC/CM Hi-C resulted in enhanced contact maps for which all metrics converged to the high signal-to-noise ratios seen in *bloomed* iHi-C (**Fig. S3**; with the exception of loop detection that only improved marginally). This highlights how *in situ* Hi-C data can also benefit from the application of *Bloom*. Together, our analyses in CMs and mESCs exemplify how *Bloom* efficiently and robustly reveals occult multi-scale interaction patterns from sparse data to identify thousands of TADs, loops, and even the more labile enhancer-promoter interactions.

### Bloom outperforms state-of-the-art methods for contact matrix analysis

To assess how *Bloom* performs in comparison to existing tools for Hi-C data processing and interaction detection, we benchmarked against eight state-of-the-art tools using all datasets analyzed here (see **Methods** for details). Briefly, we created a reference set of promoter-enhancer interactions using information from EnhancerAtlas 2.0,^31^ FANTOM5,^32^ and ENCODE.^33^ In parallel, Hi-C data were qualified as either dense or sparse. Then, given the result of each method applied to each dataset (i.e., a ranked list with an *in silico* score representing putative contacts/ loops) we generated receiver operating characteristic (ROC) and precision-recall (PR) curves for each method, as well as the respective under the curve areas, AUROC and AUPR. In all cases, *Bloom* outperformed competing methods. On sparse data especially, *Bloom* performance was strikingly superior, with a difference of ~0.17 in AUROC and ~0.28 in AUPR when compared to the next best-ranked method (**Fig. S4a,b**). Using Friedman-Nemenyi tests^34^ to assess statistical significance for each AUROC and AUPR, *Bloom* again outperformed most methods with *P*<1e^−3^, and the next best-ranked method with *P*<5e^−2^ and <1e^−2^ for AUROC and AUPR, respectively (**Fig. S4c,d**). Overall, *Bloom* appears superior with regard to the sensitivity versus specificity tradeoff. Note that we chose not to include “deep” learning approaches designed to enhance Hi-C map resolution here.^35,36^ Although efficient, such methods do not inherently identify or quantify significant contacts and, thus, fall outside the scope of this work.

### Bloom inherently quantifies interaction strength and prioritizes enhancer influence

Next, we selected three of the methods used for benchmarking due to their performance and unique formulations (**Fig. S4**) to evaluate against *Bloom* as regards interaction discovery: HiCCUPS,^15^ Fit-Hi-C,^37^ and diffHiC.^38^ HiCCUPS is a parsimonious algorithm that performs best on deeply sequenced data and robustly detects strong loops (like those by CTCF). Fit-Hi-C is tailored to “mid-range” contacts (0.05-10 Mbp); it does away with “classical” binomial modeling and uses splines instead of “binning” to capture distance information and explore numeric instability learned from iterative correction matrix balancing.^39^ Finally, diffHiC is the most sensitive of the three and typically reports the most chromatin contacts; it exploits general linear modelling (GLM) and Bayesian procedures to model technical and biological variability, and performs well on shallow-sequenced data.

Using 10 kbp-resolution Hi-C maps from human umbilical vein endothelial cells (HUVECs),^4^ we focused on a well-studied 1-Mbp domain on chr8 encompassing the *MYC* locus. We retrieved all contacts deemed significant by HiCCUPS, Fit-Hi-C, diffHiC or *Bloom* and displayed them in scaled contact matrices (**Fig. 3a**). Of the five *bona fide MYC* enhancers (**Fig. 3a**, *green circles*),^40^ HiCCUPS and Fit-Hi-C retrieved only one and two, respectively. *Bloom* and diffHic retrieved all five enhancer-promoter contacts, but diffHic returns a total of 163 contacts within this 1 Mbp-long locus and appears to be “over-calling”; *Bloom* identified 11 significant contacts (**Fig. 3a**). We next focused on the 500 kbp around *MYC* using both *in situ* Hi-C^4^ and “native” iHi-C data from HUVECs^24^ to identify significant contacts in raw and *bloomed* maps. In raw *in situ* data, no interactions were called at 5 kbp-resolution, but *Bloom* allowed discovery of five enhancer-promoter and three enhancer-enhancer interactions (Fig. 3b). In unprocessed iHi-C data, one enhancer-promoter, one repressor-promoter and one unknown, but previsouly-reported, contact^41^ stood out of the sparse matrix. Following *Bloom*, essentially all known promoter-enhancer contacts were unveiled, on top of dense connectivity amongst enhancers (**Fig. 3b**). All interactions were marked by relevant epigenetic marks and DNase I hypersensitivity, as well as by strong RNAPII ChIA-PET loops (own and ENCODE data;^33,42^ **Fig. 3c**). Thus, *Bloom* significantly enhances discovery of complex, high-resolution regulatory interactions, which are generally considered more dynamic and difficult to identify in Hi-C data (compared to the prominent CTCF loop signals, for instance).

**Figure 3.**
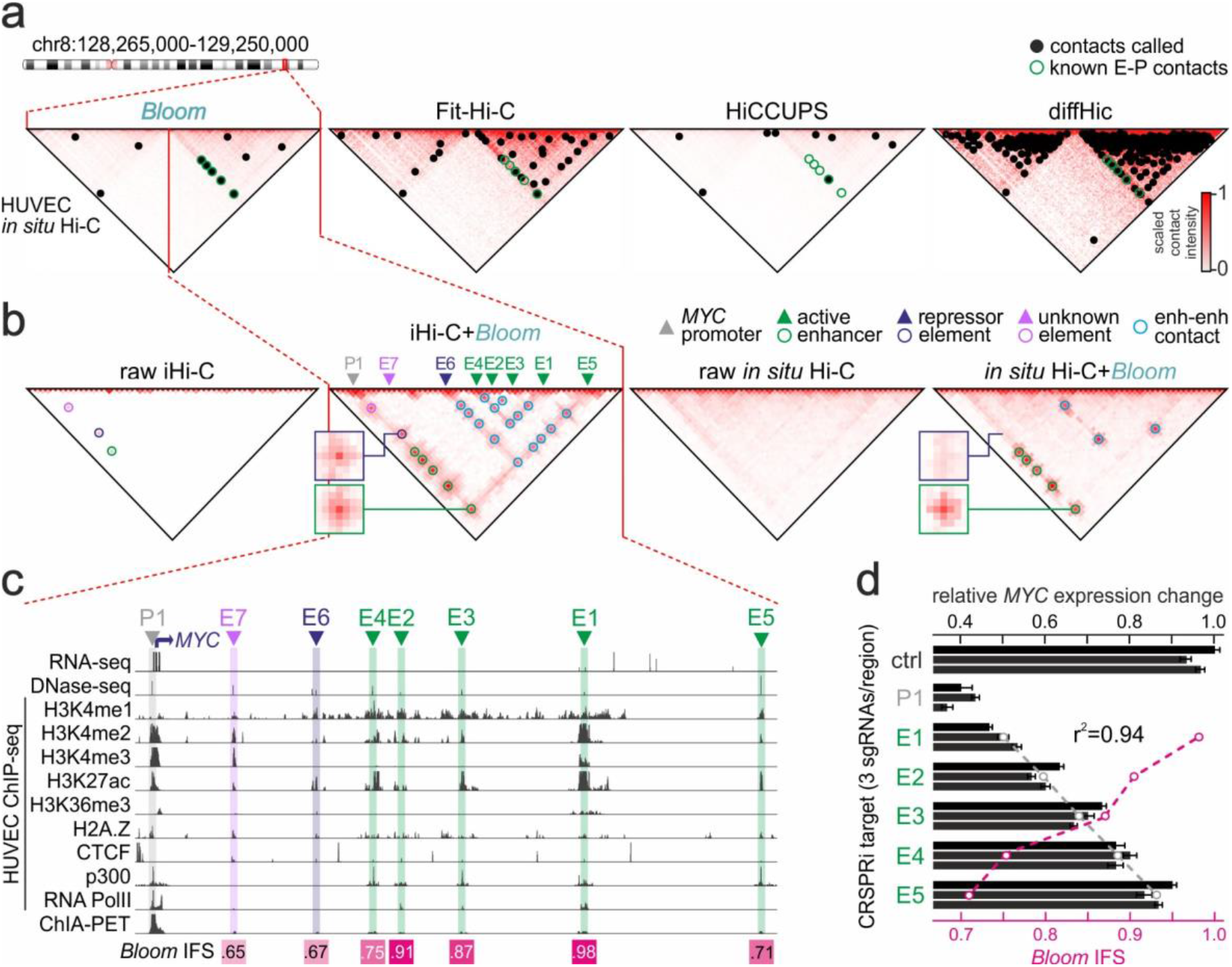
*Bloom* calls and quantifies enhancer-promoter interactions with high sensitivity. (**a**) *Bloom* contact-calling compared to three state-of-the-art methods in the 1 Mbp around *MYC* in chr8 using 10 kbp-resolution HUVEC Hi-C. Dots (*black*) show contacts called by each method; circles (*green*) denote known enhancer-*MYC* promoter contacts. (**b**) Comparison of HUVEC iHi-C and *in situ* Hi-C data in the 0.5-kbp *MYC* locus before and after *Bloom*. Called interactions (*circles*) between *cis*-regulatory elements (E1-E7) and the *MYC* promoter (P1) are denoted. Insets zoom to exemplary interaction signals. (**c**) ENCODE RNA-/DNase-/ChIP-seq and ChIA-PET data in the 0.5-Mbp *MYC* locus. *Bloom*-derived interaction strength scores (IFS) for all enhancer-promoter interactions are also shown (*bottom*). (**d**) Bar plots showing normalized *MYC* expression in three independent HUVEC CRISPRi clones (P1, promoter; E1-E5, enhancers); clones transduced with non-targeting sgRNAs provide a control (ctrl). The mean of each experimental triplet (*grey circles*) is correlated to IFS for each E-P interaction (*magenta circles*); Spearman’s correlation coefficient (r^2^) is also shown.

Given that by design *Bloom* assigns a normalized IFS value to identified contacts, we wanted to test if IFS also provides quantitative information about these regulatory interactions. The IFS values assigned to the seven key interactions between the *MYC* promoter and *cis*-regulatory elements E1-E7 ranged from 0.65-0.98 (in a 0-1 scale). We focused on the *bona fide* enhancers E1-E5, and used CRISPRi via dCAs9-KRAB to silence each individual enhancer using three different gRNAs and assessed their impact on *MYC* expression; targeting the gene promoter (P1) served as a positive control. Following RT-qPCR for *MYC* levels, we discovered an almost perfect anti-correlation between *Bloom*-assigned IFS and *MYC* suppression in our CRISPRi experiments (**Fig. 3d**; Spearman’s correlation, r^2^=0.94; *P*<10^−8^ after Benjamini-Hochberg correction). Importantly, IFS does not simply reflect the H3K27ac levels or separation of each enhancer (compare E2 to E4 in **Fig. 3c,d**) and, thus, constitutes a novel and sensitive measure of the actual influence of a given enhancer on gene activity.

### Bloom-derived IFS precisely decodes enhancer strength genome-wide

Results from the *MYC* locus suggest that *Bloom* can predict and prioritize regulatory effects in enhancer-promoter interactions from Hi-C data alone and without additional information (**Fig. 3b-d**). However, the *MYC* locus is not necessarily representative of all *cis*-regulatory scenaria across the genome. To address this, we used two enhancer perturbation datasets from K562 cells^11,12^ for which high-quality *in situ* Hi-C is also available.^4^

First, we used data from a smaller-scale CRISPRi screen, where DNase I hypersensitivity sites (HSs) within the extended loci of 7 genes (i.e. *HBG1*, *HBG2*, *PIM1*, *SMYD3*, *FADS1*, *FTH1*, and *PRKAR2B*) were targeted and gene expression changes measured.^11^ We replicated their data analysis, and used the expressed, yet unaffected, *HBE1* gene as a control. In parallel, we applied *Bloom* to K562 Hi-C to obtain IFSs for all HSs contacting genes of interest at 0.5 kbp-resolution (e,g, in the *HBG1/2*-*HBE1* locus; Fig. 4a). As before, *Bloom* produces variance-rich matrices, where differential interactions are resolved (see how HS1-4 in **Fig. 4a** specifically contact *HBG1/2* but not *HBE1*). This is not afforded by normally processed Hi-C data. Importantly, IFSs again matched closely the effects of CRISPRi perturbations. For instance, HS2 and −4 have the highest IFS values and also the strongest effect on *HBG2* expression, while *HBE1* is not at all affected (**Fig. 4b**). By extending analysis to all genes/HSs in this data, we again find a striking correlation of r^2^=0.93 between *Bloom* IFS and the screen output (*P*<10^−8^ after Benjamini-Hochberg correction). In contrast, correlation to KR-balanced Hi-C signal is only r^2^ = 0.28, and applying HiCCUPS, Fit-Hi-C or diffHiC merely increased this to 0.59, 0.52 and 0.31, respectively (**Fig. 4b**).

**Figure 4.**
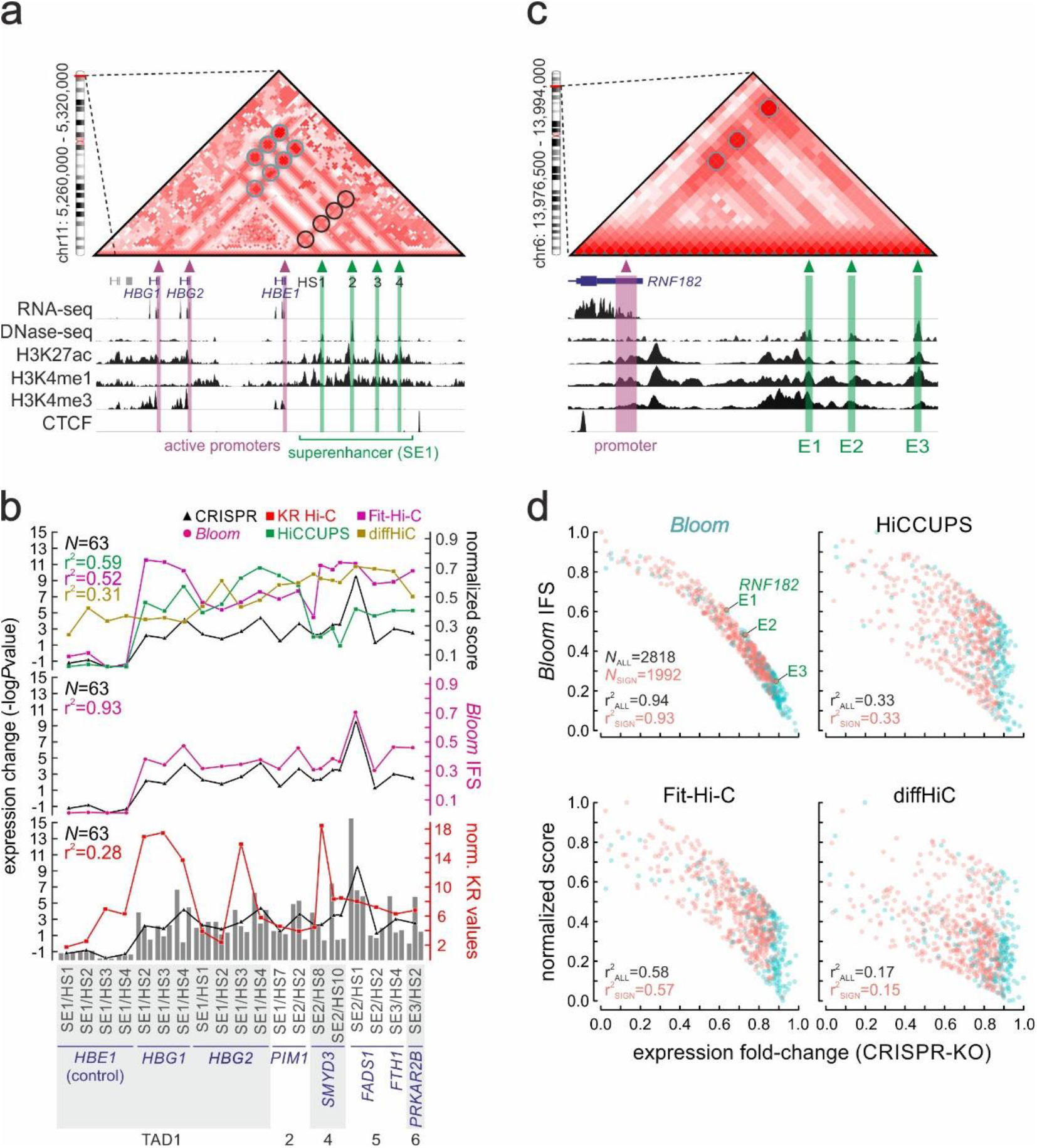
*Bloom* accurately quantifies the regulatory strength of individual enhancers genome-wide. (**a**) *Bloomed* K562 Hi-C data in the hemoglobin locus aligned to ENCODE RNA-/DNase-/ChIP-seq data. Strong (*light grey circles*) and not-called enhancer-promoter interactions (*black circles*) are denoted. (**b**) Normalized interaction frequencies calculated using four state-of-the-art methods and *Bloom* and plotted relative to expression changes after CRISPRi-targeting of multiple enhancers for seven target genes in six different TADs;^11^ expressed, but unaffected *HBE1* provides a negative control. (**c**) As in panel a, but for *RNF182*. (**d**) Scatter plots showing normalised interaction frequencies calculated using *Bloom*, HiCCUPS, Fit-Hi-C, and diffHiC relative to expression fold-changes recorded in a genome-wide screen targeting 2818 enhancers in K562 cells, 1,992 of which returned significant effects.^12^ Spearman’s correlation coefficients are shown in each case.

Next, we exploited an unbiased genome-wide CRISPR screen targeting thousands of K562 enhancers.^12^ Here, K562 Hi-C was processed as above and enhancer-promoter interactions were called genome-wide at sub-kbp resolution (see **Fig. 4c** for an example from the *RNF182* locus). Next, we plotted IFS values for each of 2,818 enhancers included in the screen or for the 1,992 that yielded significant change in the expression of their cognate gene, against the relevant fold-change in mRNA levels seen. Remarkably, regardless of using all or just “significant” enhancers, *Bloom* IFSs correlated to the screen readout with an r^2^ > 0.93 (whereas HiCCUPS, Fit-Hi-C and diffHiC gave much lower correlations of 0.33, 0.58 and 0.17, respectively; **Fig. 4d**). As this genome-wide screen covers many different regulatory scenaria across diverse chromatin loci, it conclusively verifies the ability of *Bloom* to quantify the influence conferred by individual enhacers on target promoters — and, again, this cannot be simply explained by the epigenetic features at each enhancer (compare H3K27ac and DHS signals of E1 and E3 in **Fig. 4c** with IFS/mRNA fold-change in **Fig. 4d**; for a genome-wide assessment see **Fig. S5a**). In addition, IFS does not only offer a quantitative measure for prioritizing enahncer-promoter interactions, but also high sensitivity due to its well stratified dynamic range (compared to all other methods; **Fig. S5b**).

## Discussion

Analysis tools for 3C-type data have developed alongside the advent of 3C technologies.^10,14^ However, a number of aspects in the normalization, detection, and quantification of spatial chromatin interactions embedded in Hi-C data matrices remain obscure or unresolved. We developed *Bloom* to address some of these issues. Its application to different 3C-based data matrices proved beneficial for enhancing resolution, as well as for the sensitive and reproducible detection of spatial interactions. Our main motivation came from a need to handle sparse matrices, like those derived from single-cell^22^ or “native” Hi-C experiments.^24^ We found that *Bloom* readily retrieves occult interaction patterns in such data across different scales. This is important for two reasons. First, it practically allows detection of sub-Mbp topological features in sparse data (e.g. subTADs, loops) with great accuracy and at kbp-range resolutions — something that was not feasible until now. Second, it reveals that single-cell Hi-C loops and interactions develop under the “umbrella” of an overarching and cell type-specific topology largely shared amongst individual nuclei despite the heterogeneity in gene expression and regulatory interaction rewiring.

Regulatory interactions involving gene promoters and their cognate *cis*-elements are poorly represented in Hi-C contact maps (as compared to CTCF loops, for example), and this is generally attributed to their dynamic nature. We find that contacts amongst promoters and enhancers are a prominent feature in our “native” iHi-C maps, with *Bloom* accentuating detection. In fact, in such inherently sparse data enhancer-promoter, promoter-promoter or enhancer-enhancer contacts display markedly higher signal-to-noise ratios, even when compared to deeply-sequenced *in situ* Hi-C data. Nonetheless, *Bloom* also proved useful in dense *in situ* Hi-C maps, where its application allowed enhancer-promoter contact detection with great accuracy. As a result, and given the fast implementation and moderate computational burden imposed by *Bloom*, we suggest that it can be the method of choice for analyzing not only sparse, but all kinds of 3C-type data.

Finally, *Bloom* inherently assigns a weighted interaction frequency score (IFS) to each enhancer-promoter interaction at resolutions matching those afforded by Micro-C.^20,21^ This means that one can resolve even complex enhancer-promoter communication landscapes, like those involving closely clustered “superehancers”^43^ or simply enhancers positioned very close to gene promoters, and assign stronger and weaker interactions to a given promoter. Critically, we could show, using multiple approaches pertrurbing enhancer-promoter interplay, that IFS constitutes a quantitatively accurate measure of the regulatory influence that any given enhancer confers on its target gene. This uniquely allows inference of regulatory enhancer atlases for any cell type for which Hi-C data is available without the need for multi-tiered epigenomic profiling. Such profiling can of course be helpful, but enhancer decoding via *Bloom* is independent of its output. In fact, epigenomic data alone often fail to predict “true” enhancers, and even robust models for predicting enhancer functionality, like the “activity-by-contact” (ABC) model incorporating epigenomic and KR-balanced Hi-C data, display lower precision than *Bloom* (reported AUPR at 70% recall of ~0.65 for ABC compared to >0.85 for *Bloom*).^44^ In the future, by applying *Bloom* on single-cell contact maps derived from mammalian tissues, we envisage the eventual generation of quantitative enhancer atlases spanning different cell types and unmasking regulatory heterogeneity.

## Acknowledgements

We thank the Sachinidis lab for providing cardiomyocytes for iHi-C. We also thank Ho-Ryun Chung and Emmanouil Dermitzakis for critical reading of the manuscript, and Janine Altmüller and Christian Becker at the Cologne Center for Genomics for iHi-C library sequencing. This work was supported by the Deutsche Forschungsgemeinschaft via the SPP2202 Priority program (Project Nr.: 422389065) and a Basic module grant (Project Nr.: 285697699) to AP.

## Authors’ contributions

EGG and AP conceived and designed the study; EGG performed computational analyses; AM and LB generated experimental data; EGG and AP wrote the manuscript.

## Data availability

The mouse E14 ESC iHi-C and cardiomyocyte iHi-C 2.0 data were deposited in the NCBI Gene Expression Omnibus (https://www.ncbi.nlm.nih.gov/geo/) and HUVEC iHi-C data is available via the NCBI Sequence Read Archive (http://www.ncbi.nlm.nih.gov/sra/) under accession number SRP066044.

## Code availability

The code for *Bloom* is available, under a GNU General Public License: https://github.com/eggduzao/Bloom.

## Methods

### External data sources used

We used a number of existing datasets to complement our analysis and validate *Bloom* capacities. These include high-resolution *in situ* Hi-C from GM12878, HUVECs and K562 (GEO accession: GSE63525), mESCs (GSE74055), DNase Hi-C from hPSC-derived cardiomyocytes (GSE106687), single-cell Dip-C (GSE117876) and own iHi-C from HUVECs (SRP066044). Epigenomic data were retrieved from the ENCODE (https://www.encodeproject.org) and Epigenomics Roadmap repositories (http://www.roadmapepigenomics.org). Genome and gene annotations were obtained in UCSC Genome Browser (http://genome.ucsc.edu). The focused and genome-wide enhancer CRISPR screen data were taken from the relevant published papers.^11,12^ Cell-specific enhancers used to create the gold standard for methods’ comparison were obtained in EnhancerAtlas 2.0.^31^ The single cell numbering regarding Dip-C corresponds to the numbering defined in their respective GEO submission. All external data used here were obtained or mapped to the human or mouse reference genome GRCh37 (hg19) and NCBI37 (mm9), respectively.

### Computational environment

All experiments performed in this study were performed in two different cluster settings: the Regionales Rechenzentrum der Universität zu Köln (RRZK) and the Gesellschaft für wissenschaftliche Datenverarbeitung mbH Göttingen (GWDG). At both clusters, a UNIX shell interface was used. The computing partition of RRZK contained 210 × 2 Nehalem EP quad-core processors (Xeon X5550), with 2.66 GHz average frequency and 24 GB RAM on each CPU. In GWDG, two partitions were used: (i) computationally-demanding processes were executed at a partition containing 15 × 1 Broadwell Intel E5-2650 v4 (2 × 12 cores), with 2.5 GHz average frequency and a total of 64 GB RAM on each CPU; (ii) low-/medium-demanding processes were executed at a partition containing 168 × 1 Ivy-Bridge Intel E5-2670 v2 (2 × 10 Cores), with 2.2 GHz frequency and 512 GB RAM on each CPU.

### Contact matrix processing

The initial step in *Bloom* – and the peak callers used in this study – is to obtain a 3C-data contact map from raw alignments. This is not currently implemented within our workflow. Nevertheless, multiple available tools can be used, such as Juicer^15^ or Hi-C-explorer.^46^ In brief, these tools will partition the genome in a set of non-overlapping bins ℬ = {*b*_1_, …, *b*_*N*_}. Each bin represents an interval of size ***R***. Let the initial bin be 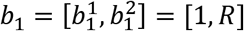, the subsequent bins can be defined as 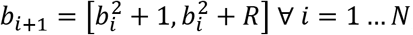; where *N* = *G*/*R* for *G* representing the size of the genome. We refer to *R* as the resolution of the experiment.

The contact map – which is the input for *Bloom* – is then created by counting the number of reads aligning to each bin. 3C approached require paired-end sequencing, where a pair of reads putatively represents one chromatin interaction event. Supposing that we have the two set of aligned reads 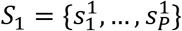 and 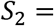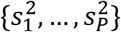. The contact map (an *N* × *N* square matrix) can be constructed as:

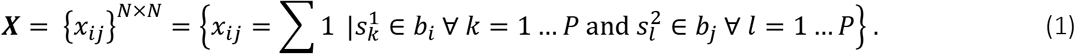

*Bloom* currently accepts all three commonly used types of contact matrix inputs: raw “sparse matrix”, “.hic” or “.(m)cool” files.

In this study, all contact maps were created using Juicer^15^ executed with default and corresponding organism-specific files, with the exception of the flags: “-r 0” (comprehensive alignment was performed) and “-t 32” (parallelization was performed using 8 processes running in 4 CPUs). With the exception of HiCCUPS – which is a part of Juicer’s pipeline – *Bloom* and all peak callers analyzed in this study, used the raw (Equation 1) or Knight-Ruiz normalized (KR)^47^ contact matrix according to their specifications and experiment design (see below).

### Bloom – sparsity verification

To define whether a matrix is sparse and should undergo SICA (see Section ***Bloom – sparsity-independent component analysis (SICA)***) or it is dense and a traditional ICA^17^ should be applied instead of SICA, we perform the following calculation.

First, we fit all values {*x*_*ij*_}^*N*×*N*^ of the original contact map’s upper matrix (∀ *i* > *j*) into a negative binomial (**Equation 1**). Then we evaluate the score 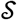 equivalent to at the 80^th^ percentile. If 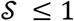, then ***X*** is sparse; otherwise ***X*** is dense. The rationale of such definition will become clearer in the following sections.

Finally, during the remainder of the algorithm, a dynamic sparsity score 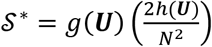, where ***U*** is the upper matrix of ***X***, *g*(***U***) is the summation of all values greater than 0 in ***U*** and ℎ(***U***) is the number of values which are greater than 0 in ***U***.

### Bloom – sparsity-independent component analysis (SICA)

We are interested in finding a set of significant vectors (“bases”) for ***X*** (Equation 1) for which we can define each contact *x*_*ij*_ as a linear combination of these bases. The Principal Component Analysis (PCA) algorithm is typically used to perform such a task, as it is able to transform the data such that variance at each “base component” is minimized.^48^ However, the “bases” – known as a low-rank matrix – are by definition orthogonal. A property that does not fully capture the complexity of 3C data.^49^ On the other hand, Independent Component Analysis (ICA)^17^ does not include orthogonality constraints. Under the principle that each contact matrix consists of a “mix” of different signals, ICA “base components” can be thought of as different “channels”, each of which captures a particular “base signal”. Nevertheless, ICA strongly relies on the fact that the “bases” – known as high-rank matrix – are non-Gaussian.^50^ Such an assumption is not possible due to the intrinsic Gaussian nature of 3C data, which is easily verified in 3C-data aggregate plots (for ax example see Fig. 2k).

Therefore, to combine the strengths of PCA and ICA we developed a methodology to retrieve a low-rank matrix using an ICA approach that is able to handle Gaussian components. Formally, let ***C*** be the covariance matrix of ***X***. Given that we are able to calculate ***C***, it will be regarded as the input to our method and can be defined in terms of the contact matrix’s bases {*p*_*ij*_}^*N*×*N*^ whose *i*^th^ row we write as:

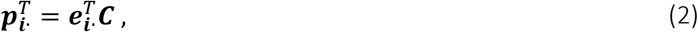

where ∙^*T*^ is the transposition operation and {*e*_*ij*_}^*N*×*N*^ = *E*[*X*]*E*[*X*]^*T*^ is the cross-expectation matrix.

To recover the columns of our independent bases, ***Y***, one at a time, consider the following procedure which takes as input ***C*** and a pair of indices (*i*_1_, *i*_2_), where *c*_*ij*_ < 0 is a randomly chosen 5-bins off-diagonal entry of ***C*** (to avoid the high entries along the contact map diagonal), and outputs column ***y***_***j***_. Note the (unknown) supports of columns and rows of ***Y*** by:

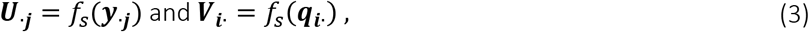

where *f*_*s*_(*v*) = {*i*|*v*_*i*_ ≠ 0}. Equation 3 implies that 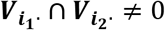. Equivalently, *i*_1_, *i*_2_ ∈ ***U***_∙***j***_ for some *j*. Using this, we typically have only one such *j*; thus, such a procedure outputs ȳ = ±***y***_∙***j***_. Then, we proceed in two steps on input ***C*** and (*i*_1_, *i*_2_).

First, we aim at finding a subset ***S*** ⊂ ***U***_∙***j***_ that contains a large fraction of the unknown support ***U***_∙***j***_. Assume ***O*** is a matrix where *i*_*ij*_ = 0 ∀ *i*, *j*, and write ***C*** as the sum of (unknown) rank-one matrices as:

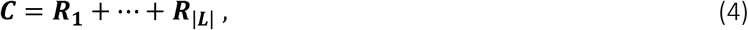

where 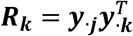 for *k* ∈ ***L***_***j*∙**_. Note that *f*_*s*_(***R***_***j***_) = ***U***_∙***j***_ × ***U***_∙***j***_. Then, the matrix 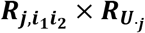 is a dense matrix of rank one. The supports of the matrices ***R***_1_ ± ··· ± ***R***_|***i***_ have small overlaps – a property preserved when adding contributions from the |***L***| − **1** terms in Equation 4. Hence, 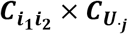 overlaps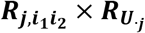 on all but a small number of entries. Therefore, 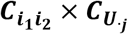 can be made to have rank one by removing the columns where these entries appear. Finally, let ***S*** ⊂ ***U***_∙***j***_ is a set of indices such that 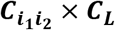 is a rank one fully dense matrix. Thus, we can evaluate the subset ***S*** ⊂ ***U***_∙***j***_ by taking the largest set ***S*** such that 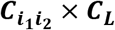 is fully dense and of rank one.

Second, given that 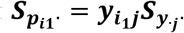, the identity 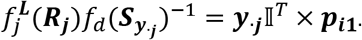 is known and 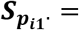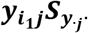, so such an identity can be written as:

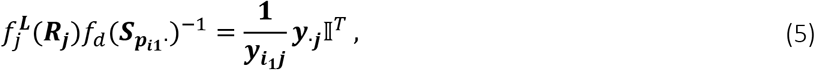

where 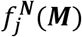 is the submatrix ***M*** with columns subset by the indexes of ***N***, and *f*_*d*_(***M***) is the diagonal vector of *f*_*s*_(*v*). The right-hand side of Equation 5 correspond to |***S***| copies of 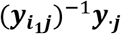 side by side. Therefore, if 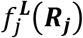 were known, ***y***_∙***j***_ would be easily calculated by taking any column of Equation 5. In the case of a symmetric input, replacing ***R***_***j***_ by ***C*** changes only a small fraction of the entries in each row. In this case, each row of:

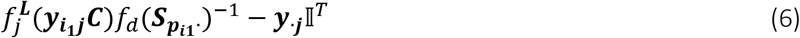

is sparse. Moreover, both 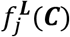 and *f*_*d*_(***S***_***pi*1∙**_)^−1^ are known, which makes the computation of 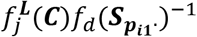 feasible. Finally, we are able to compute 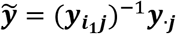, since its *i*^th^ entry is repeated several times in the *i*^th^ row of 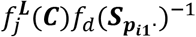. We identify 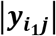 based on the fact that 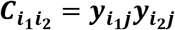 when 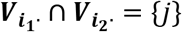, which completes the output 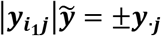. Thus, letting *f*_*m*_(*v*) and *f*_*c*_(*v*) be the mode and the median, respectively, of *v*; the final algorithm to construct 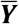 the base-normalized version of contact map ***X*** – can be computed using the following algorithm:

**Algorithm 1:**
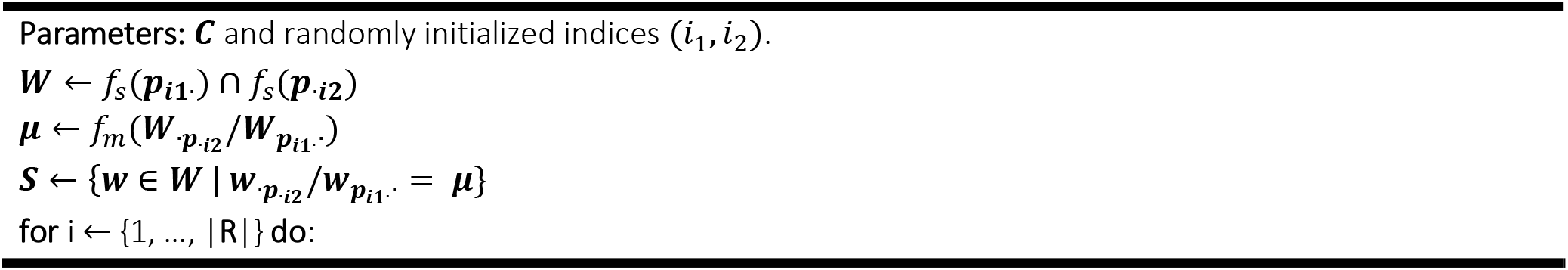
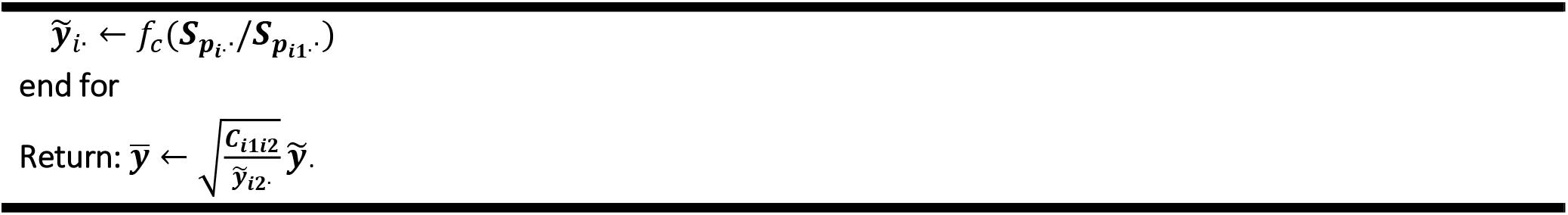
Sparse independent component analysis (SICA)

### Bloom – fast and generalized Osborne-balancing algorithm (GOBA)

The output contact map 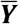 (Algorithm 1) from SICA recovers significant contacts from the raw contact map ***X*** and also handles the sparsity present in ***X***. However, as we intend, in *Bloom’*s final step, to use a sthocastic process which relies heavily, in terms of computational complexity, on how rows and columns of contact map are “linked” amongst themselves, we need to further reduce the norm in contact map ***Y***. This procedure – called matrix balancing – improves the computation of a number of features needed for the iDPMM (including the ability to parallelize massive-scale matrix multiplications; thus, enhancing execution speed; see below).^19^ We modified a well-characterized matrix balancing method called “Osborne’s balancing algorithm”^18^ by accepting a “nearly balanced” output. This is done by relaxing the algorithm’s stopping criteria aiming at considerably lower execution times.^48^ In fact, we show that this algorithm is virtually as fast as the state-of-the-art “Knight-Ruiz” method,^47^ providing a balanced matrix with no more than 2.5% instability (Fig. S6). This novel approach, termed “Generalized Osborne-balancing algorithm” (GOBA) is detailed below.

Algorithm 2 shows the original Osborne-balancing approach, which balances a matrix in the 2-norm. Importantly, this algorithm assumes that the matrix is irreducible. A matrix is considered reducible when there is at least one permutation matrix, ***M***, such that:

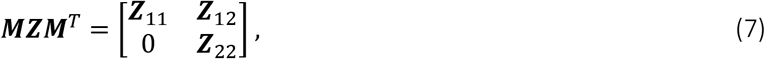

where ***Z***_11_ and ***Z***_22_ are square matrices. A diagonally similarity transformation can make the off-diagonal block, ***Z***_12_, arbitrarily small with 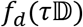 for arbitrarily large *τ*. For convergence of the balancing algorithm, it is necessary for the elements of 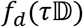 to be bound. For reducible matrices, the diagonal blocks can be independently balanced. However, a reducibility assessment of a contact map would siginificantly increase the computational time. Given that the chance of generating a reducible 3C-data contact map ***Z*** = {***Z***_*ij*_}^*N*×*N*^ is very low 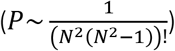, we may safely assume a contact map represents an irreducible matrix.

**Algorithm 2:**
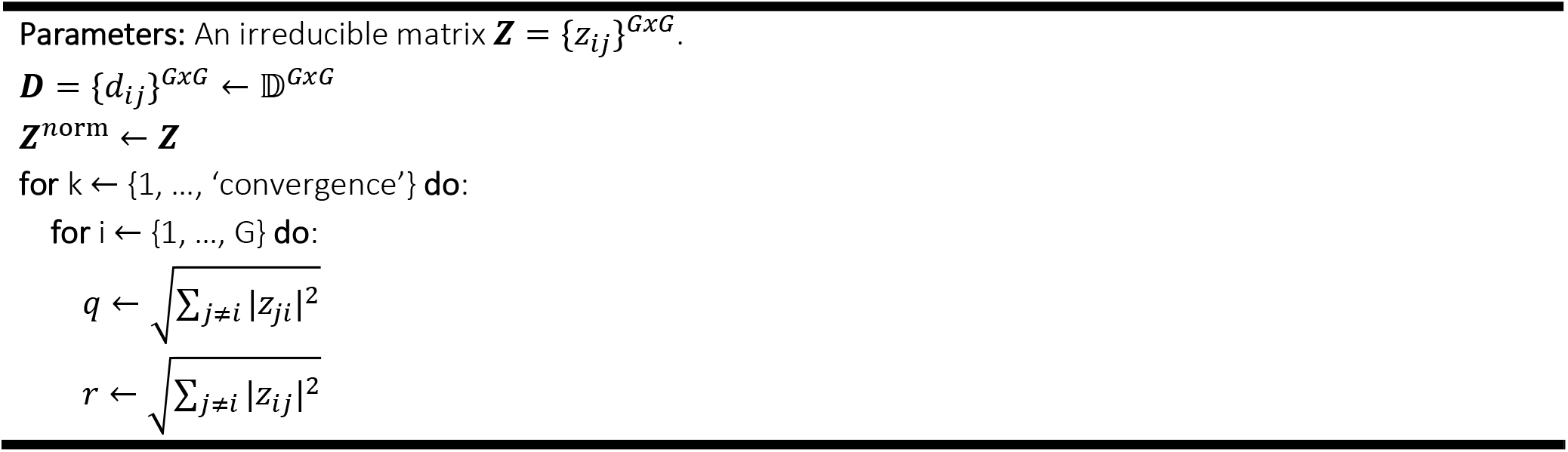
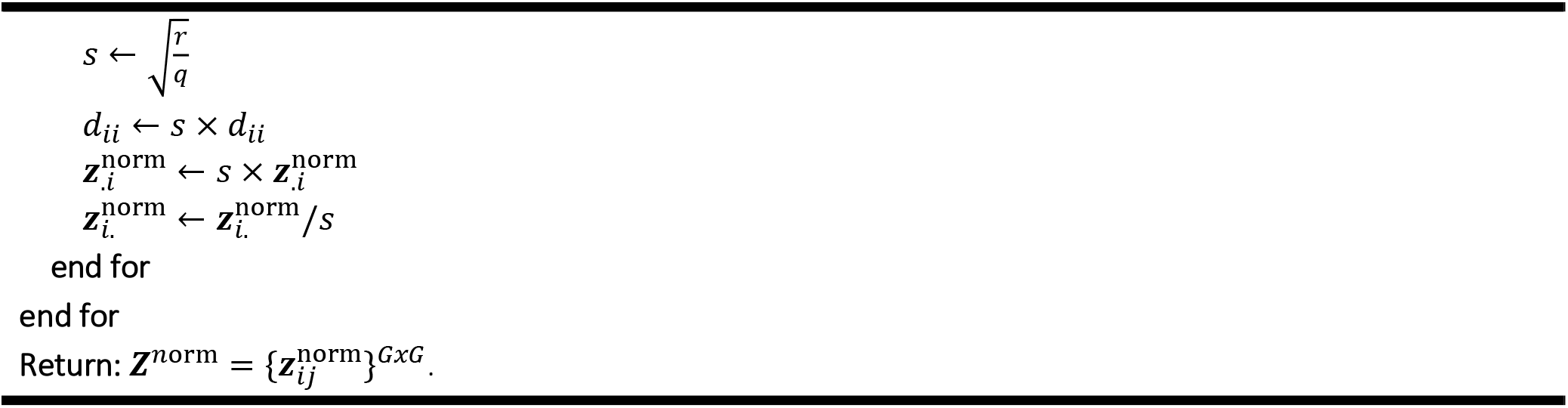
Osborne matrix balancing

This balancing algorithm acts cyclically on columns and rows of ***Z***. The quantity *s*^2^*q*^2^ + *r*^2^/*s*^2^ is minimized at 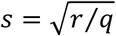, and has a minimum value of 2_*rq*_. The Frobenius norm, denoted as 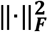 is the obvious choice for this particular case since:

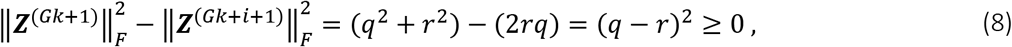

Is ever-increasing. Osborne showed that Algorithm 2 converges to a unique balanced matrix and that ***D*** converges to a unique (up to a scalar multiple) non-singular diagonal matrix,^48^ which has minimal Frobenius norm among all diagonal similarity transformations. Parlett and Reinsch generalized the balancing algorithm to any *n*-norm^51^ by changing the *q* in Algorithm 2 and *r* in terms calculation as follows:

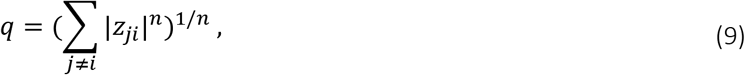

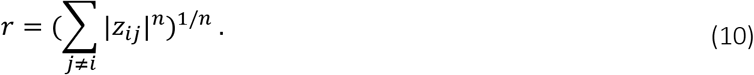

Parlett and Reinsch, however, do not quantify if the norm of the matrix will be minimized. The norm will in fact be minimized, only if, for the balanced matrix, ***Z***^***n*orm**^

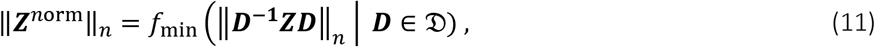

where 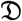 is the set of all non-singular diagonal matrices and vectorial forms of the columns of ***Z*** into a *G*^2^ column-vector. Furthermore, the diagonal element of ***D*** is restricted to powers of the radix base (typically 2), to ensure computational stability.^48^ Such an algorithm provides only an approximately balanced matrix. Doing so – without computational error – is desirable, as an exact balanced matrix is not required for our purpose. Algorithm 3 shows the orginal Parlett and Reinsch algorithm for any *n*-norm:

**Algorithm 3:**
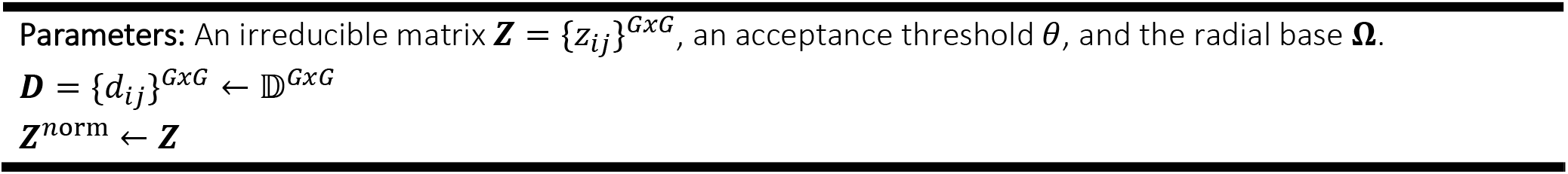
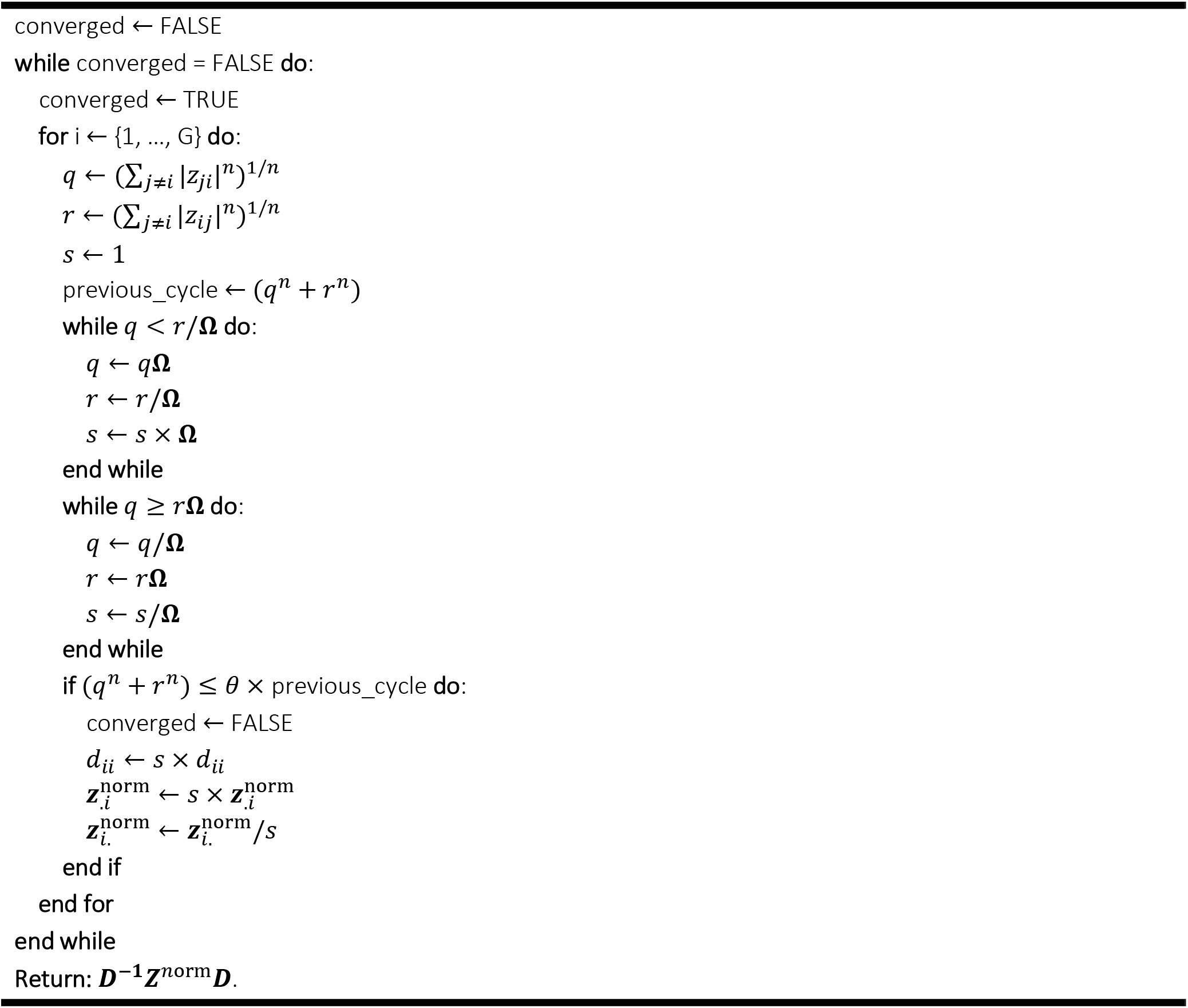
Parlett-Reich matrix balancing

Again, the algorithm proceeds in a cyclic manner, operating at a single row/column at a time. In each step, the algorithm minimizes *s*^2^*q*^2^ + *r*^2^/*s*^2^. The exact minimum is obtained for 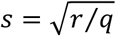. At the completion of the second inner loop, *s* satisfies the inequality:

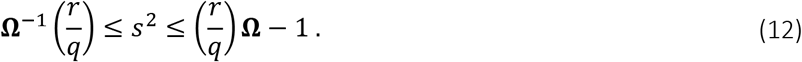

The stopping criterion states that the current step must decrease *s*^2^*q*^2^ + *r*^2^/*s*^2^ to at least *θ* of the previous (*q*^*n*^ + *r*^*n*^) value. Thus, the algorithm terminates when a complete cycle does not provide any significant decrease. Nevertheless, we observed that such a stopping criterion is not a sufficient indicator of whether the contact map’s norm truly decreases. Since the diagonal (and neighboring) elements of the current row/column is sufficiently larger than the rest of the entries, the balancing is unable to minimize the norm by a significant amount (since the diagonal element would remain unchanged). Thus, including the diagonal (and neighboring) elements in the stopping criterion provides a better indication of the relative decrease in norm. This can be acomplished by explicitly adding the diagonal-proximal elements 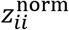 into the calculation of *q* and *r*, as

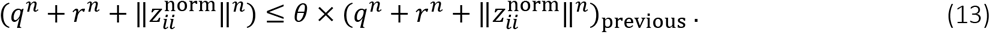

Thus, the final version of GOBA, considering as input our contact map ***Y*** and as output a GOBA-balanced matrix ***Z***^***n*orm**^, can be formalized as follows:

**Algorithm 4:**
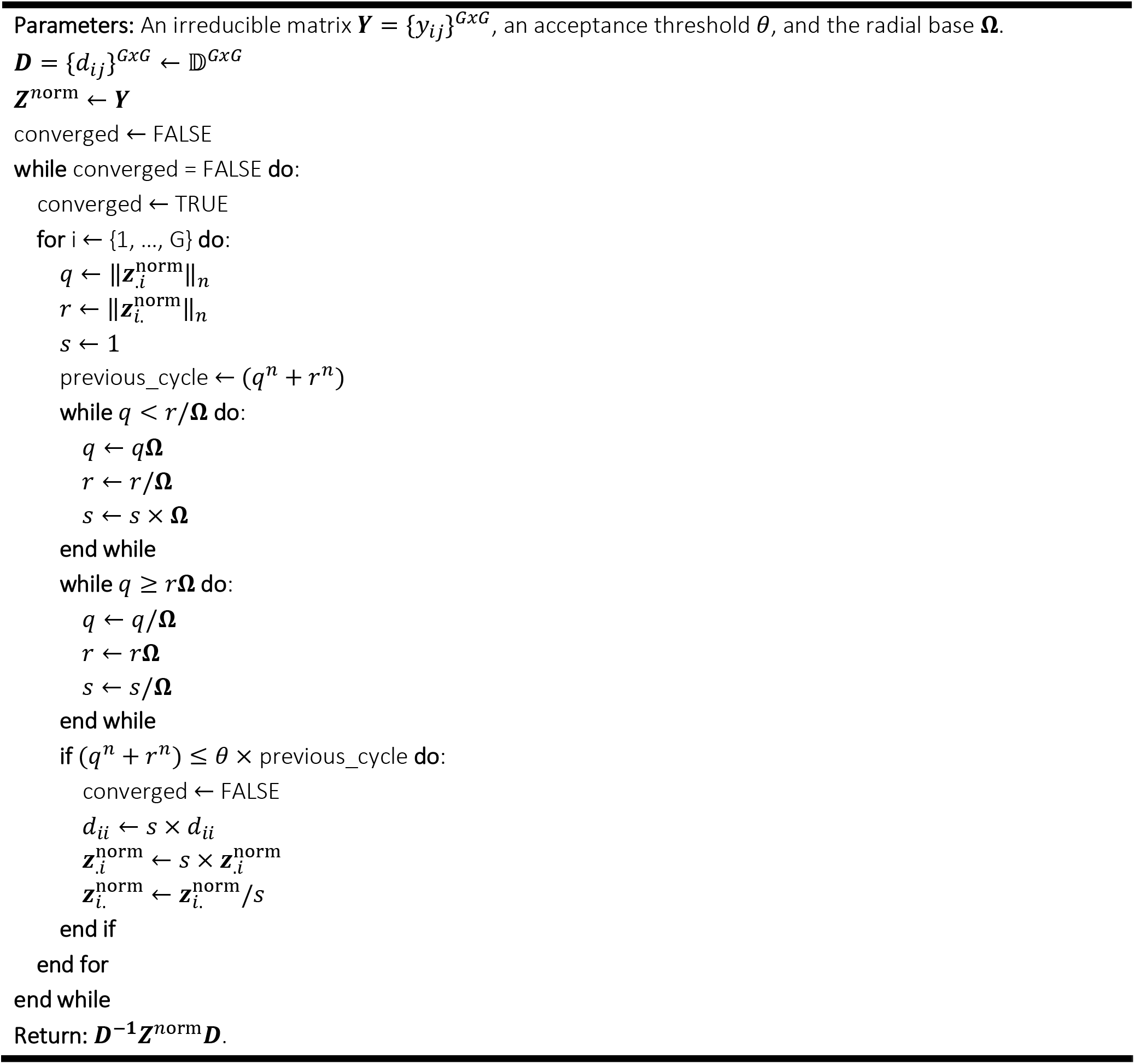
Fast and generalized Osborne-balancing algorithm (GOBA)

We have set *θ* = 0.975 within our framework (see Fig.S1b). However, we have implemented it as an open parameter; *θ* can be [0, 1]. Nevertheless, it is recommended to set *θ* ≥ 0.9 for an efficiently balanced output.

### Bloom – iterative hierarchical Dirichlet process mixture model (iDPMM)

In the final step of *Bloom* an iterative hierarchical Dirichlet Process Mixture Model (iDPMM) is applied to the contact map ***Z***^***n*orm**^; i.e. after application of SICA/ICA and GOBA. iDPMM is a nonparametric Bayesian approach for modeling grouped data, in which each group is associated with a mixture model; with final aim of “linking” these mixture models.^19^ By analogy to Dirichlet process (DP) – denoted as 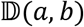 – a nonparametric prior is defined (the so-called hierarchical DP) and then used in a groupped mixture model setting.^52^ The purpose of iDPMM applied to ***Z***^***n*orm**^ is to: (i) detect and enhance significant chromatin contacts; (ii) increase contact map resolution in a binning-free manner, and (ii) automatically assign a distribution-free metric to significant contacts (which we call “interaction frequency score” – IFC) by performing a convolution of certain parameters of the Dirichlet instances as shown below. Here, we present iDPMM using the “stick-breaking construction”, which is more frequent in the literature.^19,52,53^ More formally, a hierarchical DP is a distribution over a set of random and independent probability distributions. First, we provide a formalization for the simple instance of two sets of random probability measures denoted as 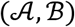. The process defines a set of random probability measures 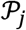, one for each group, and a global random probability measure 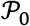. The global measure 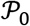 is distributed as a DP with concentration parameter 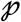 and base probability measure 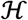 such that:

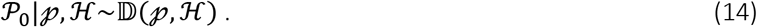

The random measures 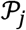 are conditionally independent given 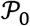, with distributions given by a DP with base probability measure 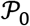:

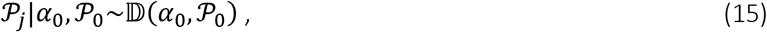

The hyperparameters of a hierarchical DP consist of the standard (baseline) probability measure 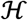 and the variation (concentration) parameters 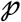 and *α*_0_. The baseline 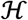 provides the prior distribution for the factors 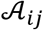. The distribution 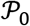 varies around the prior 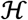, with the amount of variability governed by 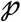. The actual distribution 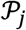 over the factors in the *j*^th^ group deviates from 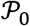, with the amount of variability governed by *α*_0_. A hierarchical DP can be used as the prior distribution over the factors for grouped data. For each *j*, let 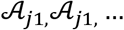 being independent and identically distributed, random variables distribute as 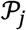. Each 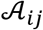 is a factor corresponding to a single observation 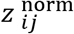. The likelihood is given by:

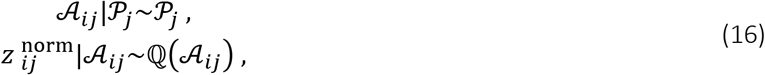

where 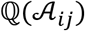 denotes the distribution of the observation 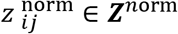 given 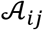, completing the definition of a hierarchical DP mixture model.

iDPMM can be extended to more than two levels by considering the baseline probability measure 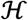 as drawn from a DP. Such a hierarchy can be enhanced for as many levels as are deemed useful. The formalization of such a concept using the “stick-breaking construction” can be performed as follows. We introduce a random set of variables *β* representing the degree of data imputation to be performed given the sparisty of the contact map (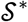; see below). Given that the global measure 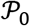 is distributed as a DP, this is expressed as:

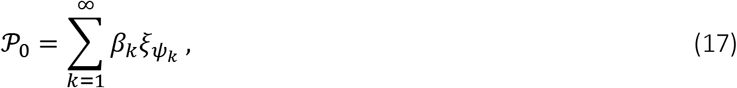

where *ξ* corresponds to a Gaussian-distributed random variable which determines the “data imputation level”; and both 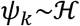 and 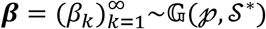 (Here 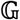 stands for the GAE – Griffiths, Engen, and McCloskey distribution)## are mutually independent. Because 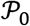 has support at the points 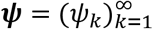, each 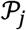 necessarily has support at these points as well, and thus can be written as:

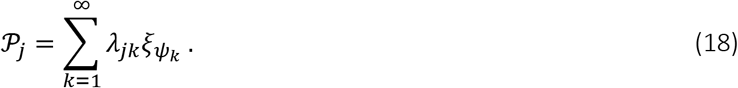

Let 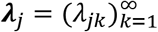. Note that the weights *λ*_*j*_ are independent given *β*, because the 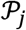 are independent given 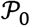. To describe the relationship between the weights *λ*_*j*_ and the global weight *β*; let {*A*_1_, …, *A*_*r*_} be a measurable partition of 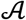 and let *K*_*l*_ = {*k* | *ψ*_*k*_ ∈ *A*_*l*_} ∀ *l* = 1, …, *r*. Note that {*K*_1_, … , *K*_*r*_} is a finite partition of the positive integers. Further, assuming that 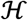 is non-atomic, the *ψ*_k_’s are distinct with probability 1; therefore, any partition of the positive integers corresponds to some partition of 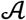. Thus, for each *j*, we have:

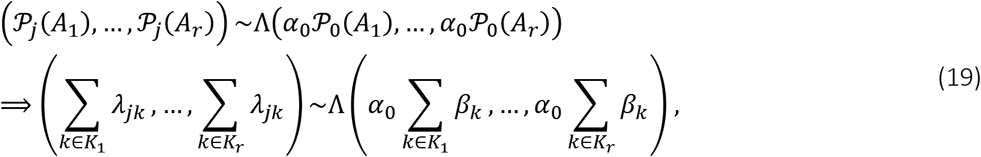

for every finite partition of the positive (weight) integers. Hence, each *λ*_*j*_ independently distributes according to 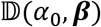, where we interpret *β* and *λ*_*j*_ as probability measures on the positive integers. If 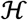 is non-atomic, then a weaker result still holds; if 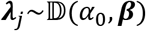, then 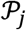 as given in Equation 18 is still 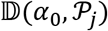 distributed.

Given that each factor *λ*_*ji*_ is DP-distributed according to 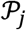, it takes on the value *ψ*_*k*_ with probability *λ*_*ji*_. Again, let *w*_*ji*_ be an indicator variable such that 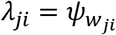. Given 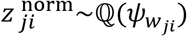. Thus, we obtain an equivalent representation of the hierarchical DP mixture through the following conditional distributions:

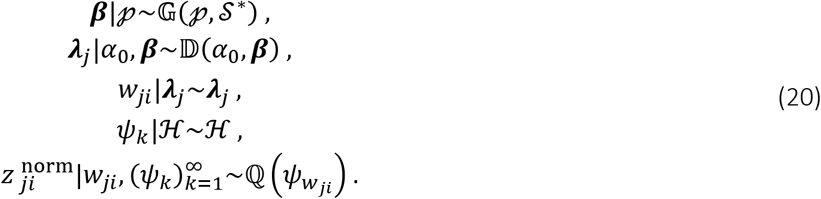

We now derive an explicit relationship between the elements of ***β*** and ***λ***_*j*_ keeping in mind that the “stick-breaking construction” for DPs defines the variables *β*_*k*_ in Equation 19 as:

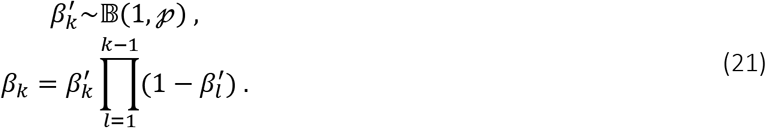

Using Equation 20, we show that the “stick-breaking construction” produces a random probability measure 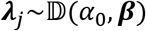:

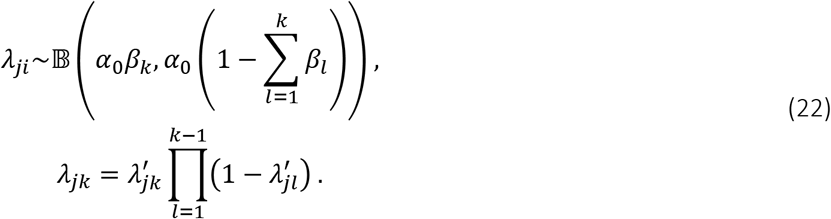

To derive Equation 22, first note that for a partition ({1, …, *k* − 1}, {*k*}, {*k* + 1, *k* + 2, …}), Equation 19 gives:

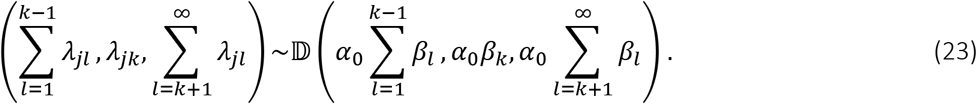

Removing the first element and using standard properties of the Dirichlet distribution,^19^ we have:

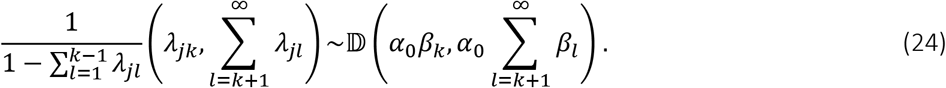

Finally, define 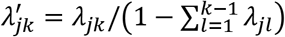 and observe that 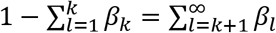. Together with Equations 17, 18 and 20, this completes the description of the “stick-breaking construction” for hierarchical DPs. *Bloom* final contact map 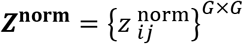, after *I* iDPMM iterations, where *G* is the reference genome size, can now be re-binned to any resolution chosen by the user. Resolutions higher than 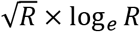, where *R* is the highest resolution that provides meaningful analyses on the original contact matrix are not recommended. In the end, all scores derived from the whole contact map are scaled to the interval [0, 1].

### Bloom – interaction frequency score (IFC)

Given our formalization of the iDPMM method, the IFC, for each instance *d* of a C-data contact map with |*D*| Dirichlet instances, can be calculated as the colvolution of all its instances. Since local contact features are already enhanced by SICA (Algorithm 1), all contacts higher than the contact matrix’s norm – given the chosen threshold *θ* in Algorithm 4 – will represent a significant contact.^19^ More formally, this calculation can be written, for a two-instances example, as:

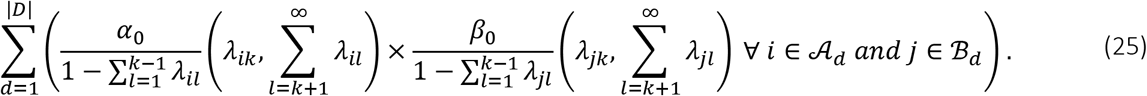

### Further contact maps processing

A/B compartments were defined using the following procedure. First, for each intra-chromosomal contact map, the eigenvector of the first principal component of the Pearson’s observed vs expected correlation matrix was determined with Juicer function “eigenvector” (v.1.19.02). Then, we calculated the average open chromatin (DNase-seq) read density for the regions in which the eigenvalues were positive (> 0) and negative (< 0). Finally, we considered as A-compartments the regions associated to the eigenvalue’s sign with the highest average DNase-seq density, and as B-compartments the remaining regions. Saddle plots were generated as previously described.^16^ Topologically associating domains (TADs) were computed using rGMAP v.1.4 on contact maps with a resolution of 10 kbp and the following parameters: logt = TRUE, dom_order = 2, maxDistInBin = 80, min_d = 25, max_d = 100, min_dp = 5, max_dp = 10, hthr = 0.99, t1thr = 0.75. Given two consecutive TADs T1 and T2, the TAD boundary strength was calculated as the division between the average of all contact map’s values inside T1 and T2 and the average contact map’s values within the same region but outside T1 and T2.

### Loop calling using Bloom and competing methods

*Bloom*’s loop calling depends on the open variables throughout the entire framework. Thus, all open variables were selected based on the average which maximizes the area under the ROC curve (AUROC) and the area under the precision recall curve (AUPR) for each individual execution on all datasets used in this study (see Fig. S6a). Such selected parameters do not maximize of the accuracy of any individual dataset; however, they enhance the ease of usage of *Bloom*, since the user does not need to perform a parameter search.^55^ Furthermore, we have observed that the accuracy between optimal dataset parameters and the global selected parameters do not significantly impact *Bloom* accuracy (*P*<0.01; Mann-Whitney-Wilcoxon test with Benjamini-Hochberg correction). On the other hand, to provide a challenging scenario for *Bloom*, all other competing loop callers used dataset-specific optimum parameters. For that, each competing method’s open parameters were tested using a grid-search and the parameter configuration that maximizes the AUROC and AUPR were selected.^55^

### Comparison between loop callers

To compare *Bloom*’s IFS ability as a loop caler, we have compared it against the following loop callers: HiCCUPS^15^, Fit-Hi-C,^37^ Fit-Hi-C 2.0,^56^ SIP,^57^ HiCPlus,^58^ diffHiC,^38^ HiCExplorer,^59^ and GOTHiC.^60^ First, we have created a *gold standard* dataset of promoter-enhancer interactions for each one of the cell types used in this study, categorized as dense: *in situ* Hi-C GM12878 and HUVEC,^4^ *in situ* Hi-C mESCs,^27^ and DNase-HiC CMs;^30^ and sparse: Dip-C GM12878,^22^ iHi-C mESCs (this study), iHi-C HUVECs,^24^ and iHi-C 2.0 CMs (this study). The intent of such a gold standard dataset is to be representative of a subset of possible interactions ocurring in a cell-specific manner. The rationale is that all methods are compared against the same gold standard, which avoids a bias towards a specific method. To create such a dataset, we have obtained all cell-specific promoter-enhancer interactions from the EnhancerAtlas 2.0. Then, we obtained the complete human enhanceosome from FANTOM5 and merged to the enhancers from EnhancerAtlas 2.0. The enhancers from FANTOM5 which are not present in the EnhancerAtlas 2.0 will constitute our “false interactions”. The interaction between FANTOM5 enhancers and promoters were created on the basis of proximity, with a strict limit of 500 kbp. Subsequently, we have obtained cell-specific RNA-seq and removed all interactions containing genes that had an expression level lower than the 90^th^ percentile of that particular cell’s expression distribution. Moreover, we obtained cell-specific DNase-seq and H3K4me1/H3K27ac ChIP-seq from ENCODE. Then, we proceed to label our interaction as follows. Valid (true) interactions consisted of all pairs of regions between a valid gene (given the expression filtering) and an enhancer from EnhancerAtlas 2.0 containing at least 50% overlap between both a H3K27ac peak and either an H3K4me1 or a DNase-seq peak. Invalid (false) interactions consisted of all pairs of regions between a valid gene and an enhancer from FANTOM5-only which did not overlap a H3K27ac peak and either a DNase-seq or H3K4me1 peak. All other possible interactions were considered as “poised enhancers”, “polycomb-repressed” or simply “ambiguous” and were removed from the final gold standard dataset. The rationale of such “strict” criteria is to create a dataset which is small enough to contain all putative loops called by the methods. This is necessary since some methods simply call loops; while others – such as *Bloom* – provide scores to every possible region. In only three cases, all regarding the dense GM12878 dataset, some methods were not able to call every possible loop: Fit-Hi-C 2.0, SIP and GOTHiC did not call 0.78%, 1.09% and 0.45% of the loops, respectively. In these cases, the lowest possible score, given each method’s metric, were assigned to non-called loops. We proceeded by creating a ranked list of all loops, per cell type category (sparse or dense) and method, in decreasing order of “loop quality”, given each method’s metric. With such a list we created the receiver operating characteristic (ROC) and precision-recall (PR) curves for each cell type category and method (see Fig. S4a,c). We also calculated the area under the ROC and PR curves, AUROC and AUPR, respectivelly. Finally, we performed a Friedman-Nemenyi test^34^ using each cell type and method’s AUROC and AUPR (Fig. S4b,d). Such a test is designed to retrieve the significance of comparison between methods using the very scenario described above.

### Correlations and further statistical tests

To account for both linearity and non-linearity, all correlations between samples were calculated based on the Spearman’s correlation coefficient (referred to as *r*). Moreover, we used the non-parametric Mann–Whitney– Wilcoxon hypothesis test to assess significance on all cases of distributions comparison, unless explicitly denoted. All tests used were two-sided and the confidence level set at 99%. The resulting *P*-values from all tests were corrected for multiple testing using the Benjamini-Hochberg approach.

### Intrinsic Hi-C (iH-C)

iHi-C was performed as described previously^24^ with the following modifications. For E14 mouse ESCs, 5-6 million cells were lifted from coated culture plates and nuclei were isolated in “physiological buffer” (PB) complemented with 8% PEG8000 and 0.4% NP-40 for 15-20 min on ice (this procedure was repeated twice to obtain intact nuclei). Next, nuclei were incubated in PB+PEG for 45 min at 30°C in the presence of 800 units *Nla*III or *Apo*I (New England Biolabs), spun, washed twice in PB+PEG, cohesive ends were filled in using biotin-dATP and the Klenow DNA polymerase subunit (New England Biolabs), pelleted and washed again, before incubating at 16°C for 6 h in the presence of T4 DNA ligase (5 u/μl; Invitrogen). Ligation was stopped by the addition of proteinase K, followed by standard DNA isolation. Biotinylated DNA length was reduced to <800 bp where needed using Bioruptor sonicator (Diagenode). iHi-C 2.0 was performed on 1-2 million CMs derived from iPSC^61^ as previously described.^25^ Note that for iHi-C 2.0, *Apo*I (New England Biolabs) was the restriction of choice, and no biotinylation was needed as libraries were prepared on the basis of “easy Hi-C”.^62^ Both standard and iHi-C 2.0 libraries were generated with the help of the Cologne Center for Genomics facility (University of Cologne, Germany) and sequenced on a HiSeq4000 platform (Illumina). Details on sequencing/mapping/quality statistics can be found in Table S1.

### Enhancer validations in the MYC locus using CRISPRi

Human umbilical vein endothelial cells (HUVECs) from pooled donors were commercially obtained (Lonza) and grown in EBM complete medium (Lonza) with supplements and 3% FBS as per manufsacturer’s instructions. Note that after transduction with tetracycline-inducible Cas9 lentiviral vectors, HUVECs were maintained in the same medium but using tetracycline-free FBS (Clontech). Lentiviral particles used for transductions were produced as follows. Approx. 700,000 HUVEC cells were plated on 6-well plates and transfected with 1 μg dVPR, 300 ng VSVG, and 1.2 μg transfer plasmid using XtremeGene9 (Roche) after 36 hours. Media was changed to DMEM with 20% HIFBS at 16 h post-transfection, and viral supernatants were harvested and filtered through a 0.45 μM syringe filter directly before use at 48 h post-transfection.

For the constructs, sgRNAs against the five *MYC* enhancers (E1-E5) and its promoter (P1; see Fig. 3c) were designed against all possible targeting sites carrying an NGG protospacer-adjacent motif (PAM), having a length of 18-24 nt, and falling within DNase I-hypersensitive footprints. Then, those with specificity scores <20 were discarded.^63^ In the end, 18 sgRNAs were selected, 3 non-overlapping per each target site, and as a negative control, 5 non-targeting sgRNAs were used (see Table S2). Next, sgRNAs were cloned into the sgOpti backbone (Addgene #71409). In parallel, the KRAB-dCas9 constructs were modified to replace the the SFFV promoter with a TRE3G promoter and the mCherry open reading frame with an IRES-GFP or IRES-BFP cassette in pHR-SFFV-KRAB-dCas9-P2AmCherry (Addgene #60954). Stable HUVEC lines expressing sgRNAs were generated using 8 μg/ml polybrene and centrifuging at 1400 × *g* for 35 min. Cells were selected for transduction after 36 h with 1 μg/ml puromycin (Gibco) for 72 h, and then maintained long-term in 0.2 μg/ml puromycin. For each target, three independent polyclonal populations were generated via infections in triplicates. Finally, each stable HUVEC line was plated at 200,000 cells/ml in 0.5 μg/ml doxycycline and harvested cells 24 h later in Trizol (Thermo) to isolate total RNA via Direct-Zol columns (Zymo Research). AffinityScript RTase (Agilent Technologies) and random primers were used to convert RNA to cDNA, and qPCR was performed using the SYBR Green I Master Mix (Roche) using the oligonucleotides listed in Table S3 as primers. RT-qPCRs were performed in triplicates for each of the three populations derived after targeting any of the six targets regions in the *MYC* locus.

## Supplemental Figures and Tables

**Figure S1.**
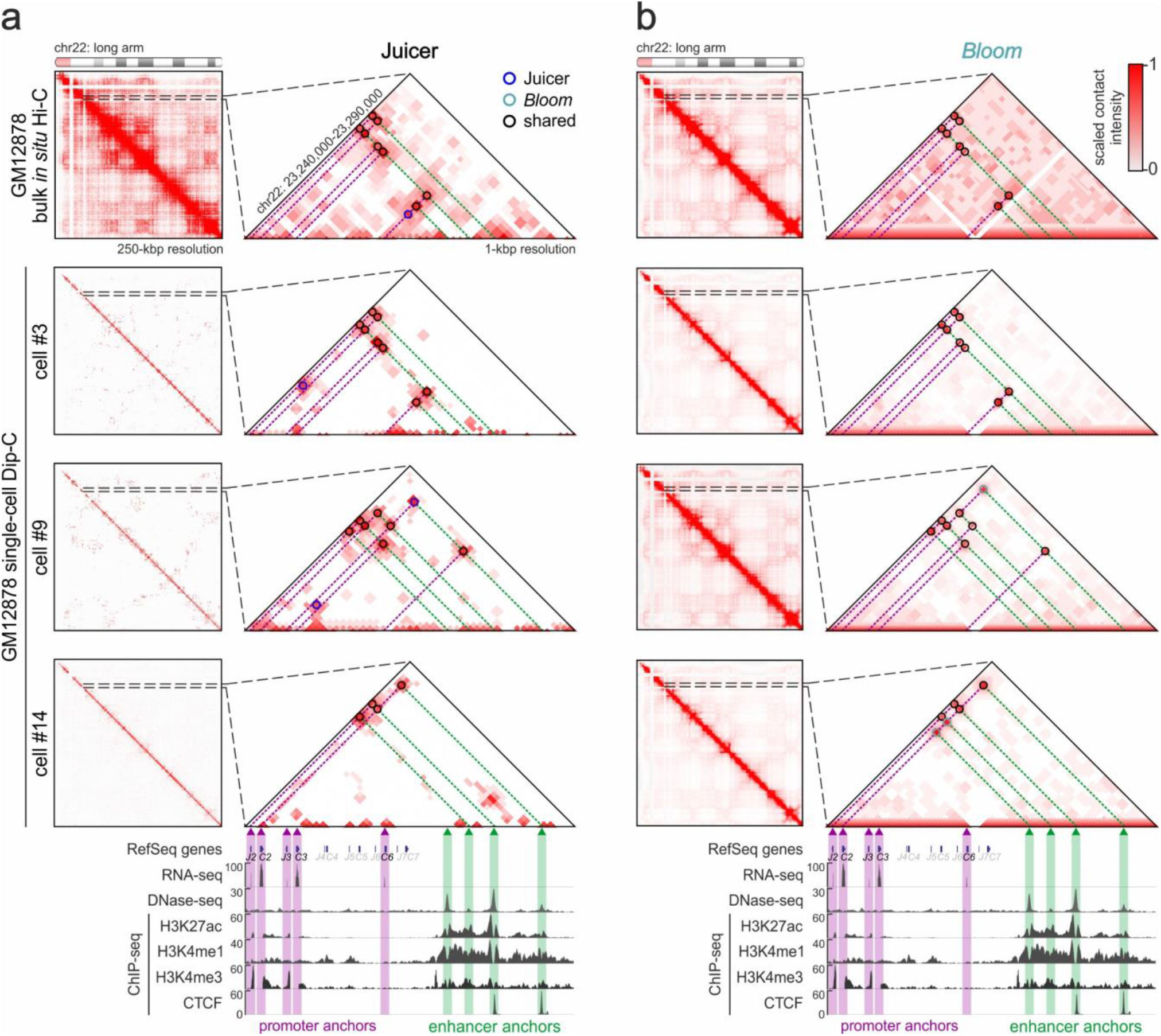
*Bloom* retrieves rich interaction profiles from single-cell Hi-C data in a gene-dense locus. (**a**) Bulk *in situ* Hi-C (*top row*) and single-cell Dip-C data from three randomly-selected GM12878 cells (#3, #9 and #14) analyzed using Juicer. *Zoom-in*: Interactions in the *IGLC/J* locus aligned to ENCODE data and all enhancer-promoter loops identified (*circles*). (**b**) As in panel b, but analyzed using *Bloom*.

**Figure S2.**
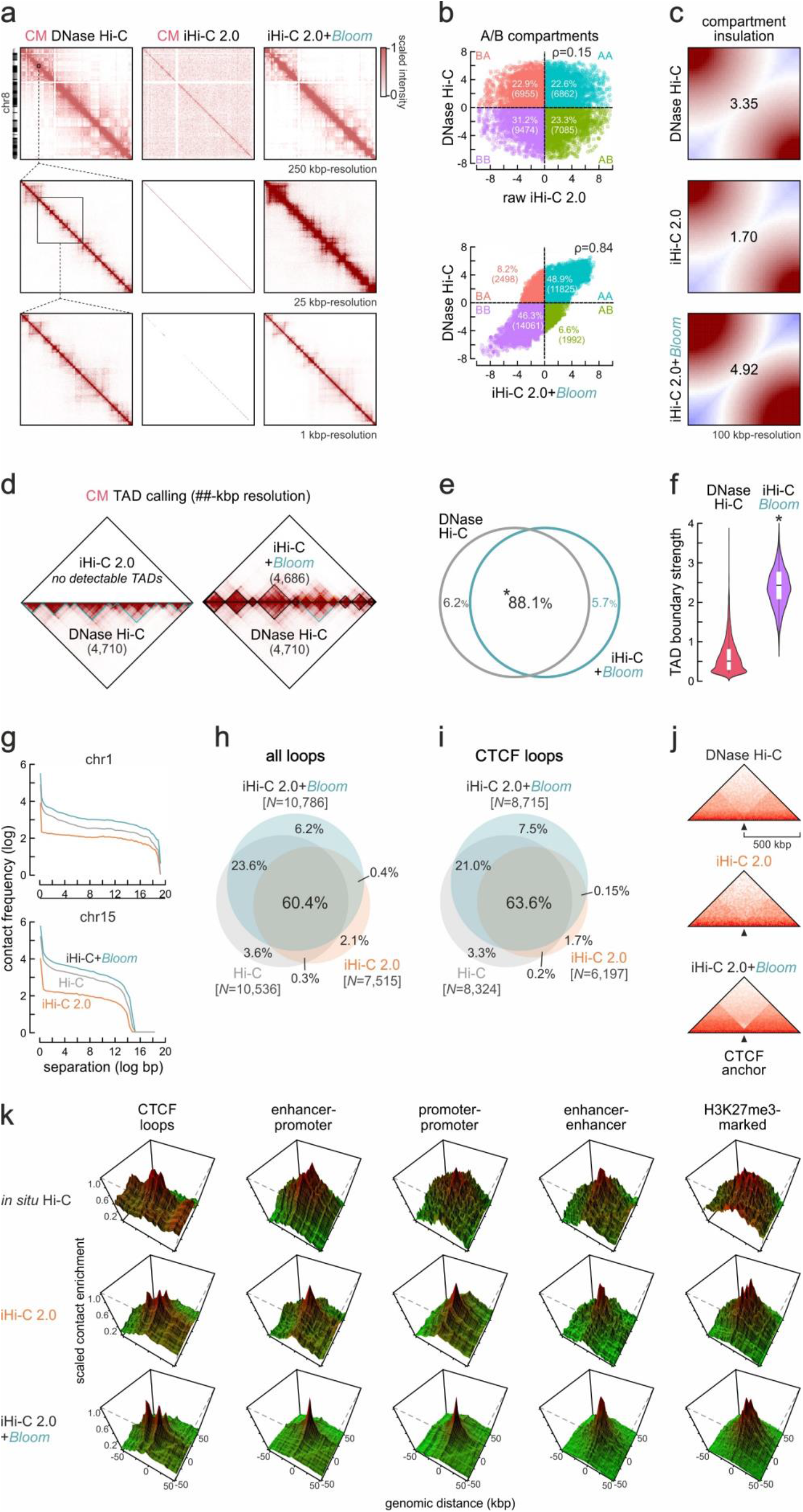
*Bloom* retrieves rich interaction profiles from iHi-C 2.0 data. (**a**) Exemplary contact maps from chr8 using DNase Hi-C (*left*), raw (*middle*) or *bloomed* iHi-C 2.0 data from iPSC-derived cardiomyocytes (CM; *right*). (**b**) Scatter plots of A/B compartment values derived using the datasets in panel a. (**c**) Saddle plots showing insulation of A-/B-compartments. (**d**) Comparison of TADs called using DNase Hi-C (*bottom triangles*) in CMs to those called using raw (*left*) or *bloomed* iHi-C 2.0 (*right*). TADs that differ in *bloomed* iHi-C data are denoted (*orange triangles*). (e) Venn diagram showing the overlap of TADs called in Hi-C (*grey*) or *bloomed* iHi-C 2.0 (*light blue*). (**f**) Violin plots showing insulation strength at the boundaries of TADs from panel d. *: significantly different; *P*<10^−5^, Wilcoxon-Mann-Whitney test. (**g**) Line plots showing contact frequency decay with distance in DNase Hi-C (*grey*), raw (*orange*) or *bloomed* iHi-C 2.0 (*light blue*). (**h**) Venn diagrams showing overlap of loops called in the three datasets from panel a. *: more than expected by chance; *P*<0.001. (**i**) As in panel h, but for CTCF-anchored loops. (**j**) Heatmaps showing mean Hi-C signal in the 1 Mbp around CTCF loop anchors from panel i. (**k**) 3D plots depicting Hi-C signal enrichment at (*from left to right*) CTCF-bound sites, active promoter-enhancer pairs, active promoters, active enhancers or H3K27me3-marked sites.

**Figure S3.**
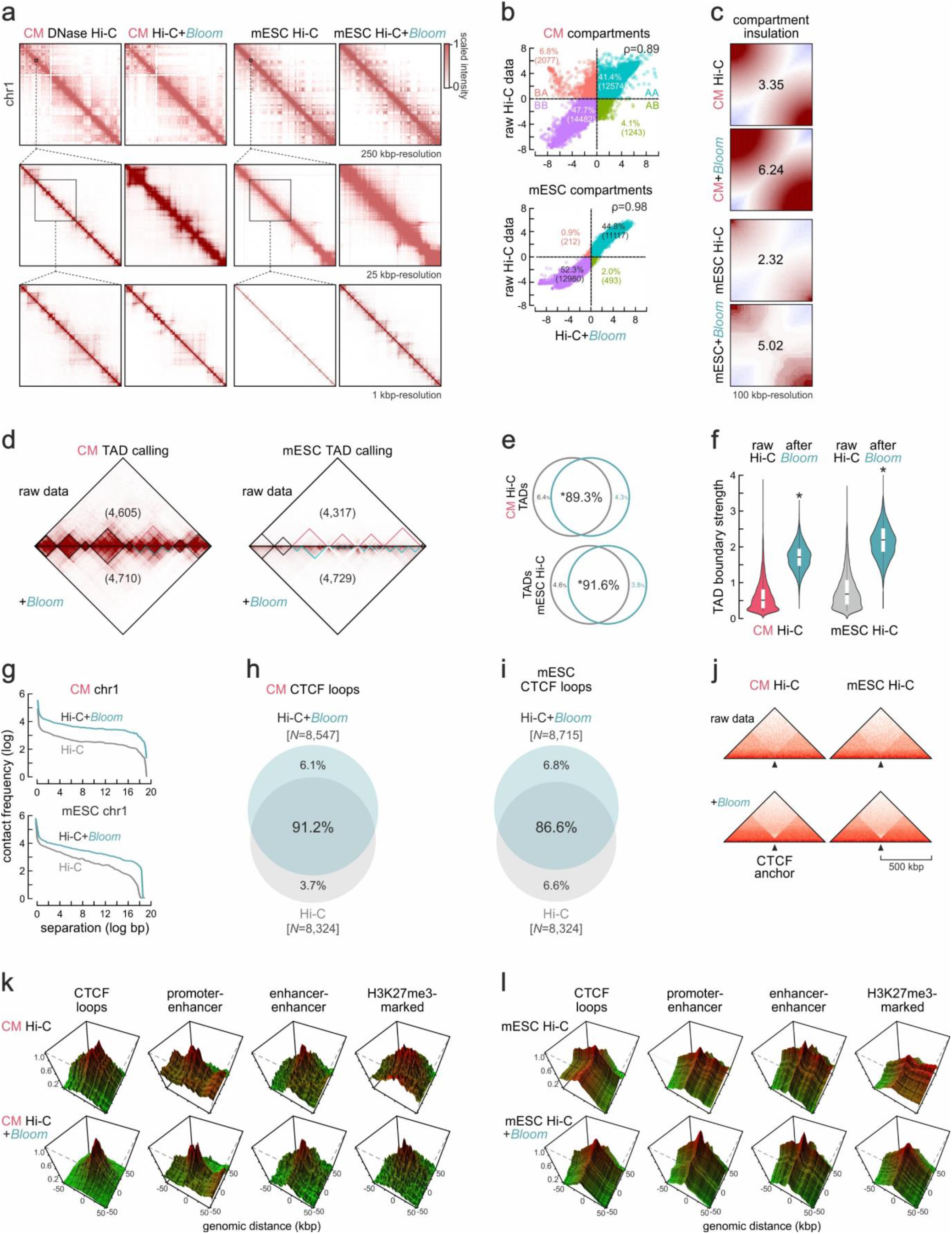
*Bloom* improves resolution and loop detection in dense Hi-C data. (**a**) Exemplary contact maps derived from cardiomyocyte (CM) DNase Hi-C (*left*) or mESC *in situ* iHi-C data (*right*) before and after *Bloom*. (**b**) Scatter plots of A/B compartment scores derived using the datasets in panel a. (**c**) Saddle plots showing insulation of A-/B-compartments for the data in panel a. (**d**) Comparison of TADs called using raw (*top*) or *bloomed* Hi-C data (*bottom*) from CMs or mESCs. TADs that differ in *bloomed* data are denoted (*light blue triangles*). (**e**) Venn diagram showing the overlap between TADs called in raw (*grey*) and *bloomed* Hi-C data (*light blue*). *: more than expected by chance; *P*<0.001. (**f**) Violin plots showing insulation scores at the TAD boundaries from panel d. *: significantly different; *P*<10^−5^, Wilcoxon-Mann-Whitney test. (**g**) Line plots showing contact frequency decay with distance in CM (*top*) or mESC Hi-C (*bottom*) before and after *Bloom*. (**h**) Venn diagrams showing overlap of CTCF loops called in CM Hi-C before and after *Bloom*. *: more than expected by chance; *P*<0.001. (**i**) As in panel h, but for mESC CTCF loops. (**j**) Heatmaps showing mean Hi-C signal in the 1 Mbp around CTCF loop anchors using data from panels h,i. (k) 3D plots depicting mean CM-specific Hi-C signal enrichment for interactions involving (*from left to right*) CTCF-bound sites, active promoter-enhancer pairs, active enhancers or H3K27me3-marked sites before and after *Bloom*. (**l**) As in panel k, but using mESC Hi-C data.

**Figure S4.**
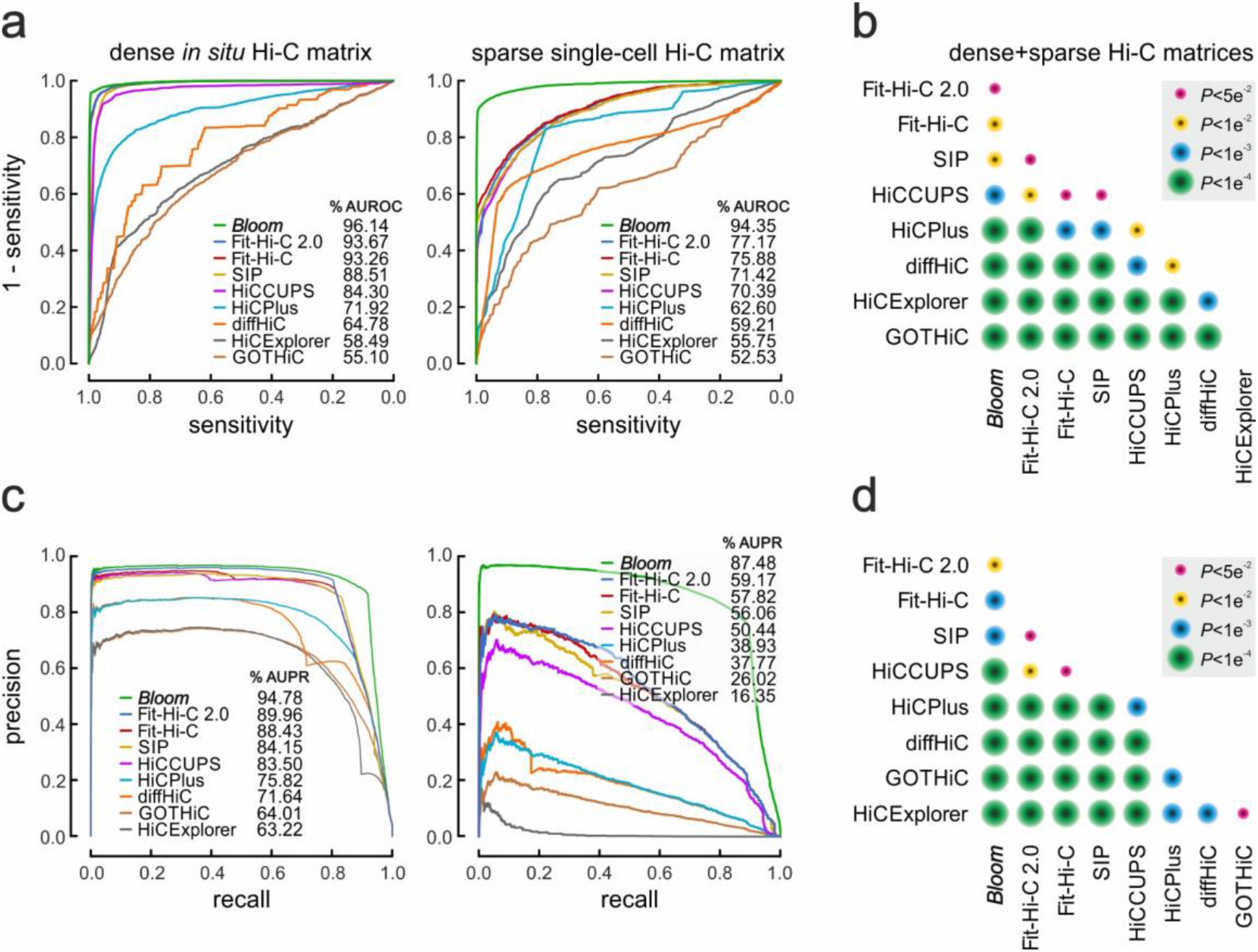
*Bloom* performance compared to state-of-the-art methods. (**a**) Receiver operating characteristic (ROC) curves for *Bloom* and another 8 computational methods on dense (GM12878 *in situ* Hi-C; *left*) or sparse matrices (Dip-C; *right*). (**b**) As in panel a, but using precision-recall (PR) curves. (**c**) Friedman-Nemenyi-testing for comparing each method’s area under the ROC curve (AUROC) across all data used; the colour code denotes how significantly the method in each column outperformed the methods in rows. (d) As in panel b, but comparing each method’s area under the PR curve (AUPR).

**Figure S5.**
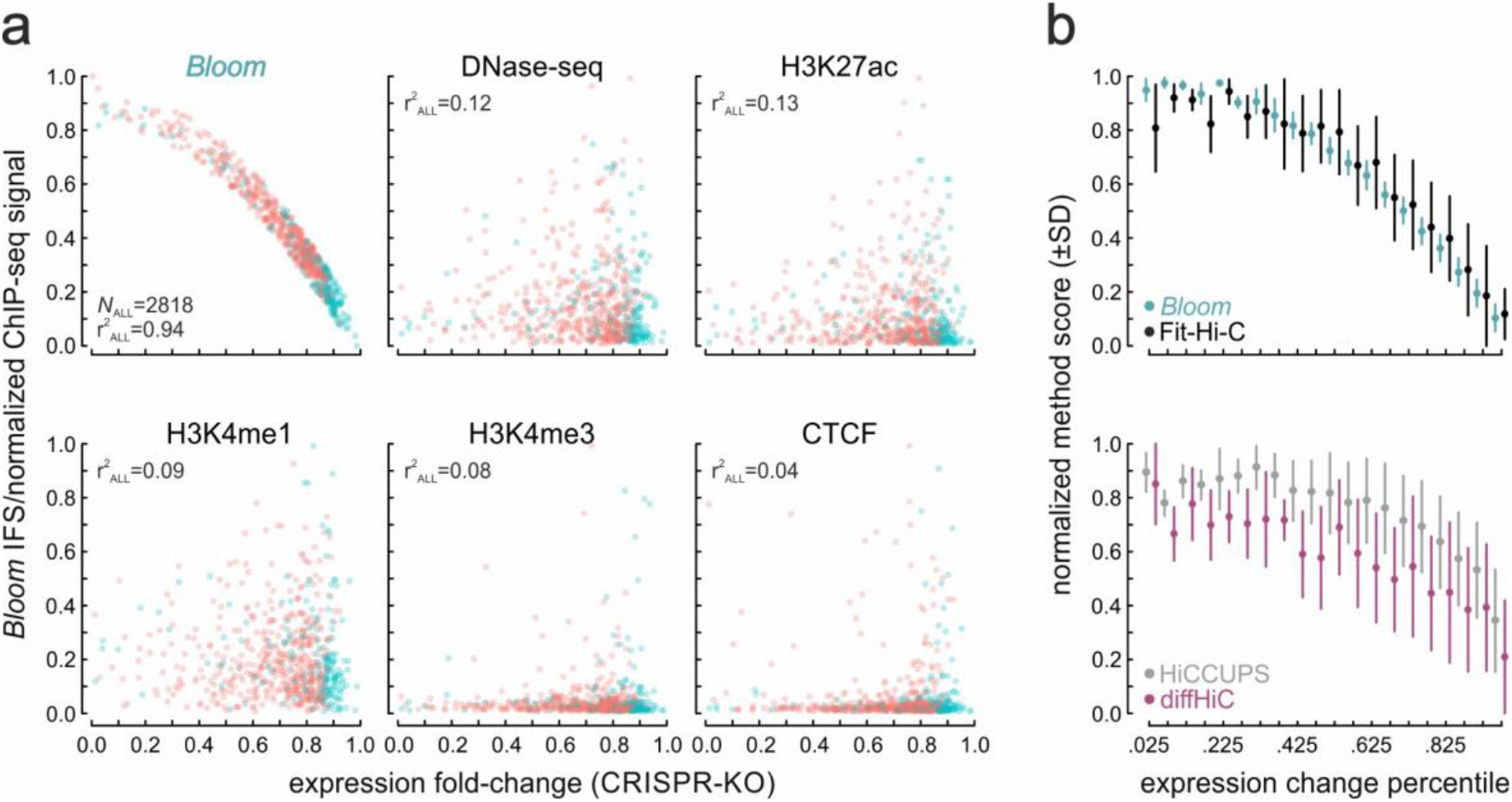
IFS quantifies enhancer strength with precision, without merely reflecting epigenetic marks. (**a**) *Bloom*-derived IFS (*top left*) or normalized DNase-/ChIP-seq signal plotted against fold-changes in gene expression from a genome-wide K562 CRISPR screen targeting 2818 enhancers; Spearman’s correlation coefficients are shown. (**b**) Normalised interaction scores (±SD) calculated using *Bloom* (*light blue*), Fit-Hi-C (*black*), HiCCUPS (*grey*) or diffHiC (*purple*) plotted over twenty consecutive expression fold-change quantiles from the CRISPR screen in panel a.

**Figure S6.**
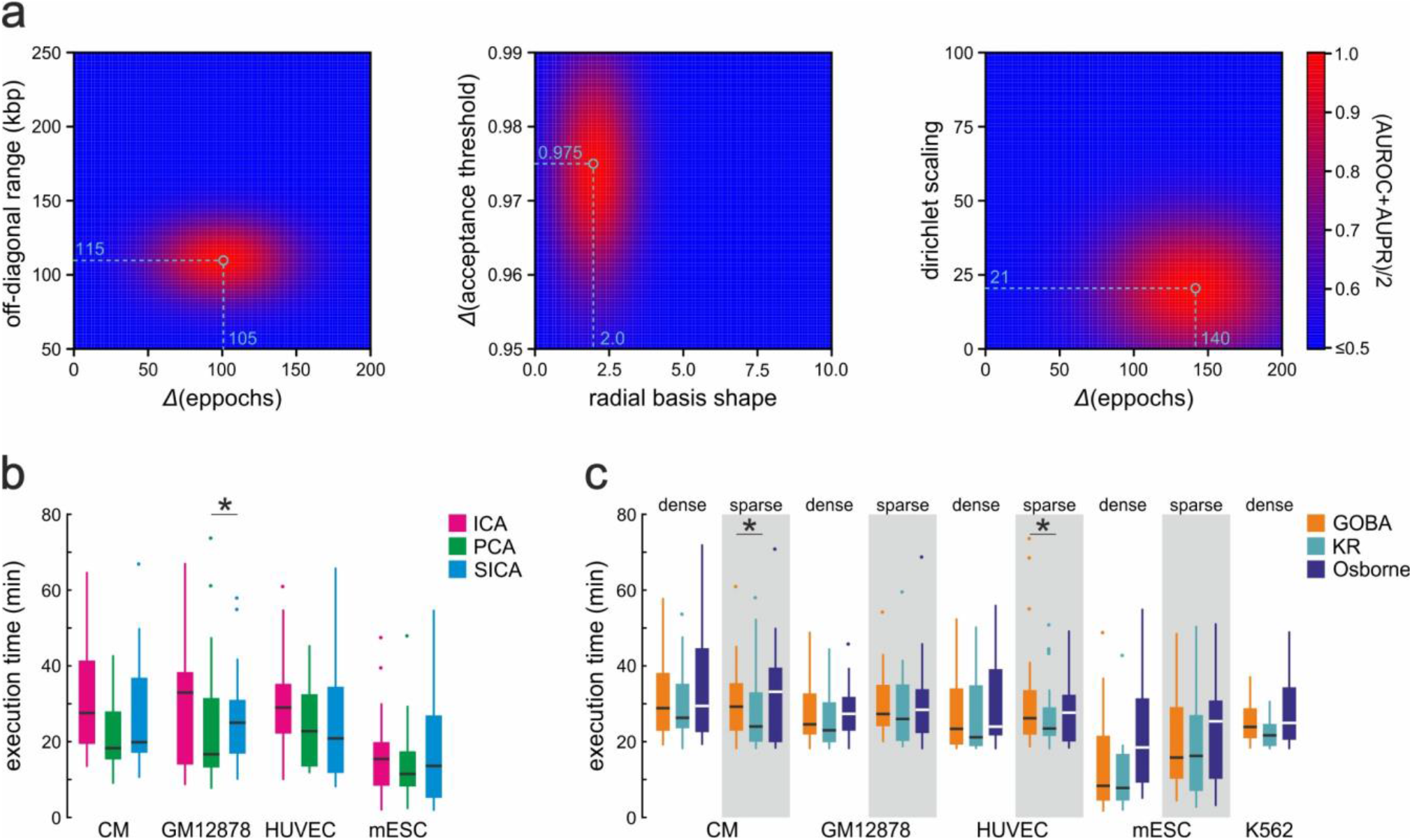
Analysis of *Bloom* algorithm’s parameterization and computing time. (**a**) *Left*: Average performance of SICA algorithm in multiple combination of its open parameters with regard to AUROC and AUPR. *Δ*Eppochs refer to the scaled difference between the gain of a current eppoch *t* versus its previous tested eppoch *t-1*. Optimal average parameters selected are shown in green. *Middle*: As in the left panel, but for GOBA parameters. *Right*: As the left panel, but for iDPMM parameters. (**b**) Computational time comparison between dimensionality reduction methods ICA (independent component analysis), PCA (principal component analysis) and our SICA (sparsity-independent component analysis) on each chromosome using the “sparse” Hi-C data indicated. *: significantly slower; *P*<0.01, Wilcoxon-Mann-Whitney test. (**c**) As in panel b, but for the matrix balancing methods Osborne, KR (Knight-Ruiz) and GOBA (generalized Osborne balancing algorithm). *: significantly slower; *P*<0.05, Wilcoxon-Mann-Whitney test.

**Table S1.**
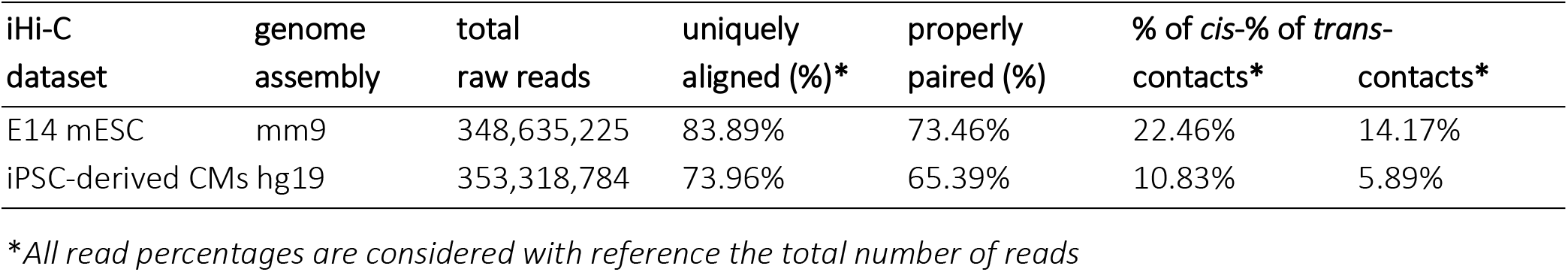
General iHi-C data statistics.

**Table S2.**
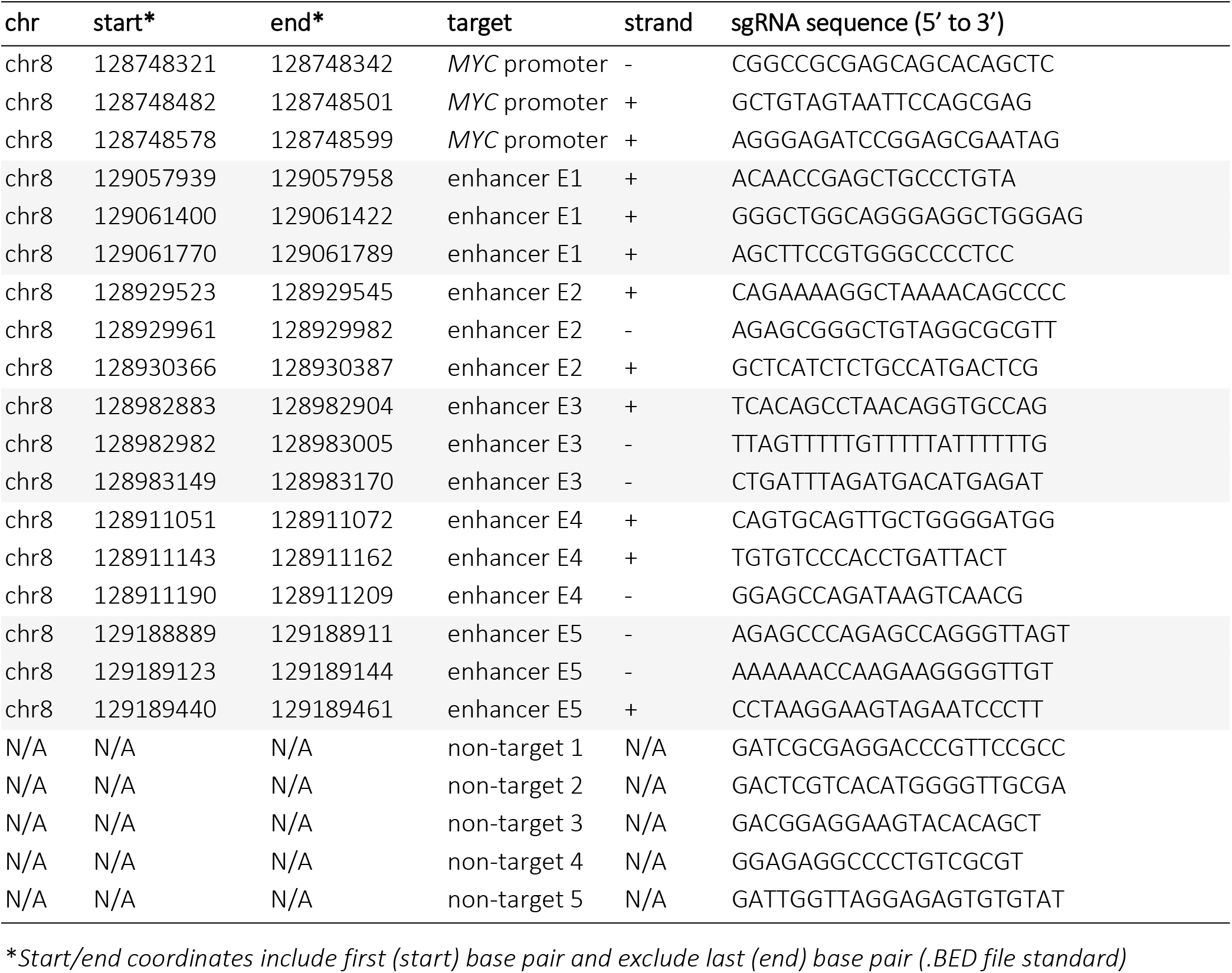
CRISPRi sgRNAs targeting the *MYC* locus.

**Table S3.**
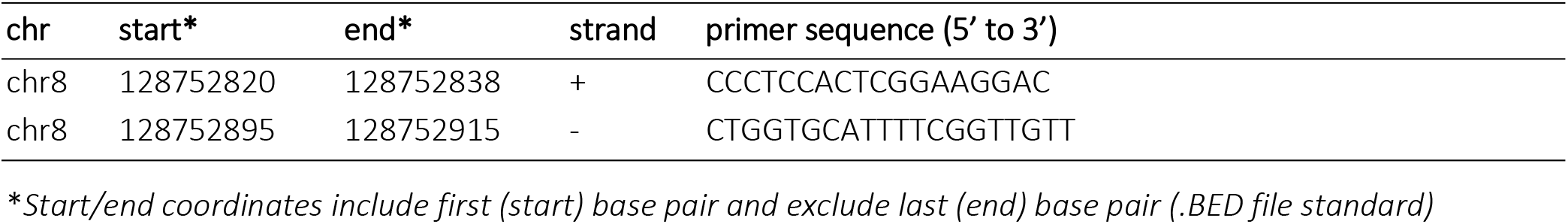
CRISPRi RT-qPCR primers for the *MYC* gene.

